# Mechanosensation Dynamically Coordinates Polar Growth and Cell Wall Assembly to Promote Cell Survival

**DOI:** 10.1101/299008

**Authors:** Valeria Davì, Hirokazu Tanimoto, Dmitry Ershov, Armin Haupt, Henry De Belly, Remi Le Borgne, Etienne Couturier, Arezki Boudaoud, Nicolas Minc

## Abstract

How growing cells cope with size expansion while ensuring mechanical integrity is not known. In walled cells, such as those of microbes and plants, growth and viability are both supported by a thin and rigid encasing cell wall (CW). We deciphered the dynamic mechanisms controlling wall surface assembly during cell growth, using a novel sub-resolution microscopy approach to monitor CW thickness in live rod-shaped fission yeast cells. We found that polar cell growth yielded wall thinning, and that thickness negatively influenced growth. Thickness at growing tips exhibited oscillating behavior with thickening phases followed by thinning phases, indicative of a delayed feedback promoting thickness homeostasis. This feedback was mediated by mechanosensing through the cell wall integrity pathway, which probes strain in the wall to adjust synthase localization and activity to surface growth. Mutants defective in thickness homeostasis lysed by rupturing the wall demonstrating its essential role for walled cell survival.

## INTRODUCTION

Growth control is of fundamental importance in biology, from the regulation of macromolecular assembly, cell physiology, up to organ development (Holley, 1975; Hong et al., 2016; Lander, 2011; Mahadevan and Mitchison, 2005). The growth of individual cells, for instance, has crucial implications for cell size determination, tissue homeostasis or cancer progression (DeBerardinis et al., 2008a; Deberardinis et al., 2008b; Fantes and Nurse, 1977; Marshall et al., 2012; Thompson, 2010). To date, however, the mechanisms which control cellular growth remain poorly understood. This is because growth is an integrated output of multiple intertwined biochemical and biomechanical elements which dynamically probe and alter the cell surface to accommodate surface expansion. Those may include surface material synthesis mediated by processes such as exocytosis and endocytosis (Hepler et al., 2001; Novick and Schekman, 1979), as well as osmotic forces and mechanical elements which set the elasticity of the cell surface, like the actin cortex, the glycocalyx or the CW (Davi and Minc, 2015; Huang and Ingber, 1999; Salbreux et al., 2012). How those modules which act at various time and length scales may dynamically feedback onto each other to control the rate of cell surface expansion remain an outstanding open question.

Cell growth has been admittedly best documented in single walled cells, such as bacteria, fungi, or pollen tubes (Harris and Theriot, 2016; Mitchison and Nurse, 1985; Qin and Yang, 2011; Rojas et al., 2014). Those are some of the fastest growing cells, with some fungal cells elongating at rates up to several tens of microns per minute (Lew, 2011; Qin and Yang, 2011). Size expansion in those cells is irreversible and limited by the synthesis of a thin and rigid sugar-made CW around the plasma membrane. The growth of the CW is thought to involve a complex balance between sugar synthesis which build the wall, and mechanical expansion driven by large internal turgor pressure which put the wall under tension to deform it (Cosgrove, 2005; Davi and Minc, 2015; Harold, 2002; McKenna et al., 2009). Because the CW bears large turgor-derived stress, it also provides mechanical integrity to those cells: removal or weakening of the CW yields near-immediate cell lysis and death. Given those considerations, cell growth may be seen as a dangerous process, as uncontrolled expansion and consequent thinning of the CW could compromise cell survival. To date, however, our understanding of how CW composition and thickness may be dynamically controlled during cell growth has been limited, as historically it has been mostly studied with Electron Microscopy, which has biased our appreciation of the CW towards a static structure.

The biochemistry and genetics which support CW growth and composition, has been extensively studied in model yeast cells (Lipke and Ovalle, 1998; Perez and Ribas, 2004). The rod-shaped fission yeast *Schizosccaromyces pombe*, for instance, serve as a prime model for walled cell tip growth, stereotypical of many bacteria, fungi and plant cells (Chang and Huang, 2014; Chang and Martin, 2009; Davi and Minc, 2015). The fission yeast CW behaves as a thin elastic shell with an elastic modulus estimated to be around 30-50MPa which resists an internal turgor pressure of 1-1.5 MPa (Abenza et al., 2015; Atilgan et al., 2015; Minc et al., 2009). In those cells, CW synthesis is restricted to cell tips during phases of cell elongation and to the cell middle for septation. At cell tips, wall synthesis is catalyzed by one α-glucan (Ags1) and three β-Glucan (Bgs1, Bgs3 and Bgs4) synthases, which elongate glucan fibers, as well as glucanosyl-transferases enzymes (Gas1 and Gas2) and one putative exo-glucanase, Exg2, which may remodel and/or crosslink glucan fibers at cell tips (Davi and Minc, 2015; Perez and Ribas, 2004). Those enzymes are under the control of the activity of the highly conserved Rho-GTPases, Rho1 and Rho2, which are activated by the GEFs Rgf1 and Rgf2 (Arellano et al., 1999; Garcia et al., 2006). Damages on the CW, such as those caused by antifungal agents are monitored by the Cell Wall Integrity pathway which can activate Rho1 and Rho2, and the expression of a set of CW repair genes through the Pmk1 MAPK (Perez and Cansado, 2010).

To understand how those different biochemical layers dynamically influence and probe the mechanics of the CW, we here introduce a novel microscopy method to directly measure CW thickness all around live growing fission yeast cells. We find that CW thickness at growing cell tips is highly dynamic in time, with phases of thickening followed by thinning phases, and *vice e versa*. We demonstrate a homeostatic system which accounts for thickness oscillations and stable values at the cell population. Homeostasis is supported by mechanosensing activities of the CWI which probe cellular growth as a mechanical stress and dynamically adjust synthesis to growth rates. Mutants defective in homeostatic thickness lyse by over-thinning the CW. This work evidence mechanochemical feedbacks promoting growth control and cell survival.

## RESULTS

### A sub-resolution method to monitor cell wall thickness dynamics in living cells

The fission yeast CW is a thin layer of typically 100-200nm, below the resolution of light-microsopy. Given large internal turgor pressure, the plasma membrane is plastered against the internal face of the CW (Osumi, 2012). In addition, glycosylated galactomannan proteins are secreted and retained at the most outer surface of the wall, and can be detected with specific lectins (Horiseberger and Rosset, 1977). We exploited those properties to develop a sub-resolution method to compute CW thickness all around single live growing fission yeast cells. We labeled the inner and outer faces of the CW respectively using an integral membrane SNARE protein tagged with a GFP at its intracellular tail (GFP-psy1) (Nakamura et al., 2001) and by adding Lectins from *Griffonia simplicifolia* that specifically bind the most outer face of the fission yeast CW, labeled with a different emitting fluorophore. Following image registration and cell contour segmentation, we computed the distance between the centers of the two fluorescent signals along lines bisecting the cell surface, which provided a local measurement of CW thickness (Chugh et al., 2017; Clark et al., 2013). Thickness measured by those means could be calculated with a precision around ~30 nm and a lateral resolution of ~500 nm, which we represented as colored cellular maps (Figure 1A and Figure S1A-S1E) (see STAR methods). Measurements were slightly sensitive to growth environment and media but robust to variations in lectin-bound fluorophores or membrane associated GFP signals (Figure S2A-S2C).

**Figure 1.**
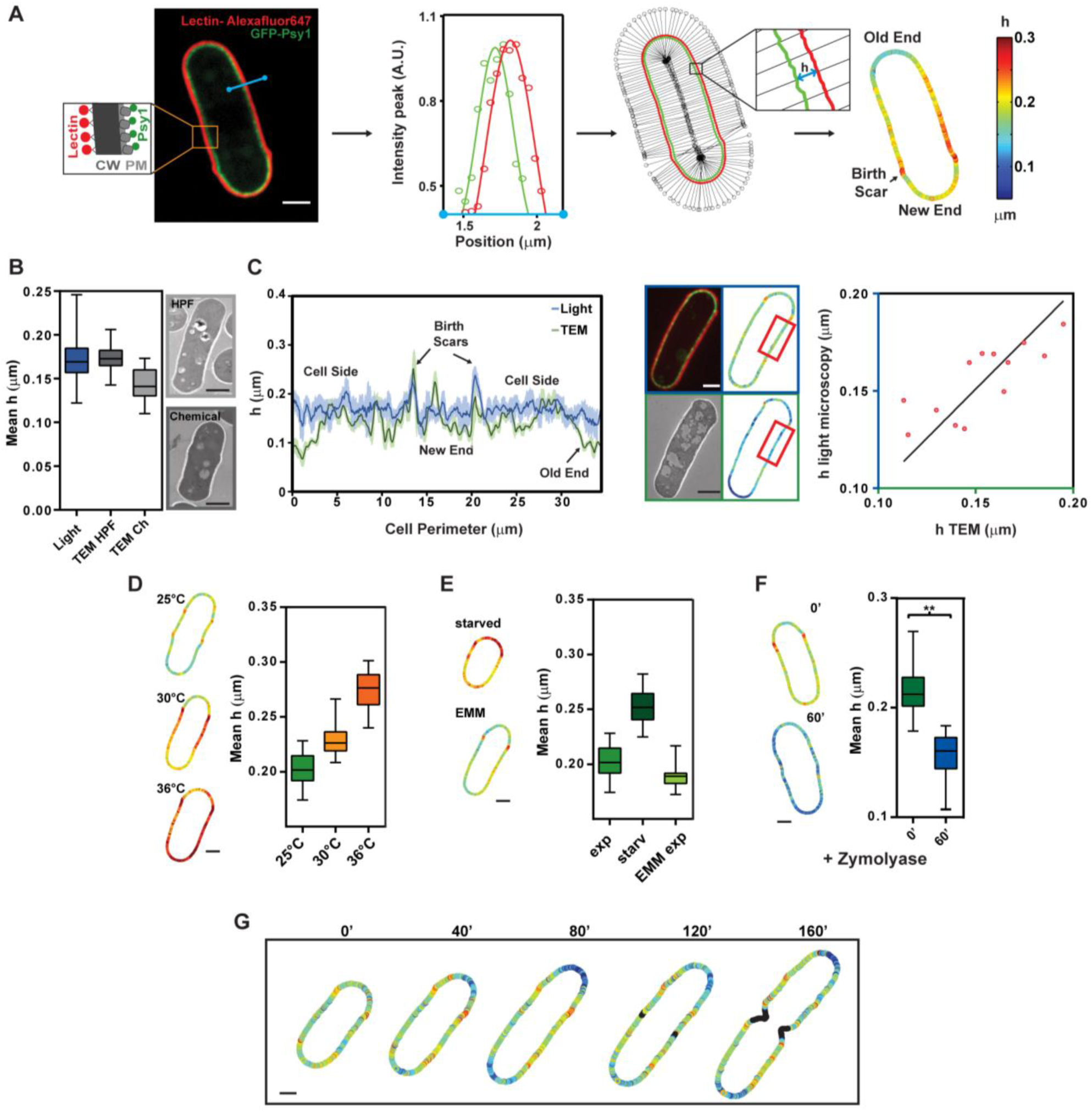
A novel method to image cell wall thickness around living cells. **(A)** Mid-slice confocal image of a fission yeast cell expressing GFP-Psy1 (plasma membrane, PM) and labeled with lectin-Alexafluor647 (cell wall outer surface). Contours are determined from the centers of Gaussian fits of each signal across the cell surface (blue line). After chromatic shift registration, the distance between the two contours yields local CW thickness measurements, h. (Right) Thickness color map around a cell. **(B)** Measurement of CW thickness using Photonic microscopy (n=41) and Transmission electron microscopy (TEM) performed by High pressure freezing (HPF) (*n=23*) or chemical fixation (Ch) (*n=22*). Representative TEM images in these conditions. **(C)** (Left) Thickness of the same single cell, shown in the middle, measured by TEM or light microscopy using Correlative Light-Electron Microscopy. (Right) Light vs TEM thickness measurement averaged over regions of ~2µm (red rectangle) (n=13 regions, from 9 cells). The line is a linear fit. **(D)** Mean wall thickness of cells grown at different temperatures, and representative thickness color maps in the same conditions (n>25 for each condition) **(E)** Mean wall thickness of cells measured in exponential phase (exp) or after ~16 h of starvation (starv), and in cells grown in minimal (EMM) medium (n>25 for each condition). **(F)** Thickness before (n=35 cells) or after 60’ of Zymolyase treatment (n=19); **P < 0.0001. **(G)** Time-lapse of CW thickness maps of a growing cell. Black points correspond to positions in which thickness cannot be measured accurately. Scale bars 2 µm; Whiskers plots show median and full data set range.

To validate our approach, we first compared it with Transmission Electron Microscopy (TEM) measurements, which serve as the current standard to image the CW (Osumi, 2012). The mean CW thickness in a wild-type population measured by our method was 170 ± 23 nm. This was comparable to values obtained from High Pressure Freezing TEM (172 ± 14 nm) and ~16% higher than values obtained in chemical fixation TEM (143 ± 19 nm), plausibly because chemical fixation may not fully protect the CW against subsequent resin embedding (Osumi, 2012) (Figure 1B). Next, we used Correlative Light Electron Microscopy to image and compute CW thickness in the same cell with TEM and our method. This yielded nearly similar patterns of thickness along the cell boundary, supporting the reliability of our method as a local measurement of CW thickness (Figure 1C and Figure S2D-S2G).

To assess the accuracy, precision and range of this live imaging method, we next tested it against conditions thought to affect CW thickness. As previously suggested, we detected a dose-dependent increase in CW thickness with temperature or as a result of prolonged starvation (Figure 1D-E) (Cassone et al., 1979). Cells treated with zymolyase, a CW digesting enzyme, exhibited significant CW thinning (Figure 1F). Finally, thickness could be imaged in the same cell over several hours with no major cellular damage and no impact of bleaching on the measurement (Figure 1G, Figure S3A-S3B and movie S1 and S2). Thus, it was possible to quantify the dynamics of CW thickness and assembly with unprecedented accuracy in populations of live growing cells.

### Polar growth sites are associated with thinner and softer cell walls

Given that wall assembly in fission yeast is restricted to growing cell tips during interphase and to the cell middle during septation, we computed CW thickness patterns in cell populations (Chang and Martin, 2009; Davi and Minc, 2015). Birth scars which are inherited from previous cell division were associated with a significantly thicker CW, with a thickness of h_scar_ = 224 ± 20nm. Interestingly, we observed that the CW was consistently thinner at the old end, the end which grows most during interphase, h_oe_ = 130 ± 26 nm, as compared to cell sides, h_side_ = 170 ± 20 nm. Thickness at the new end, which only grows moderately after New End take Off (NETO) was in contrast only slightly thinner than cell sides, h_ne_ = 167 ± 14 nm (Figure 2A). We also found that monopolar tips of outgrowing spores, spheroplasts, mating projections or of *tea1Δ* mutants branching from cell sides were also markedly thinner at the front growing site than on the non-growing back (Bonazzi et al., 2014; Kelly and Nurse, 2011; Mata and Nurse, 1997; Petersen et al., 1998) (Figure 2B).

**Figure 2.**
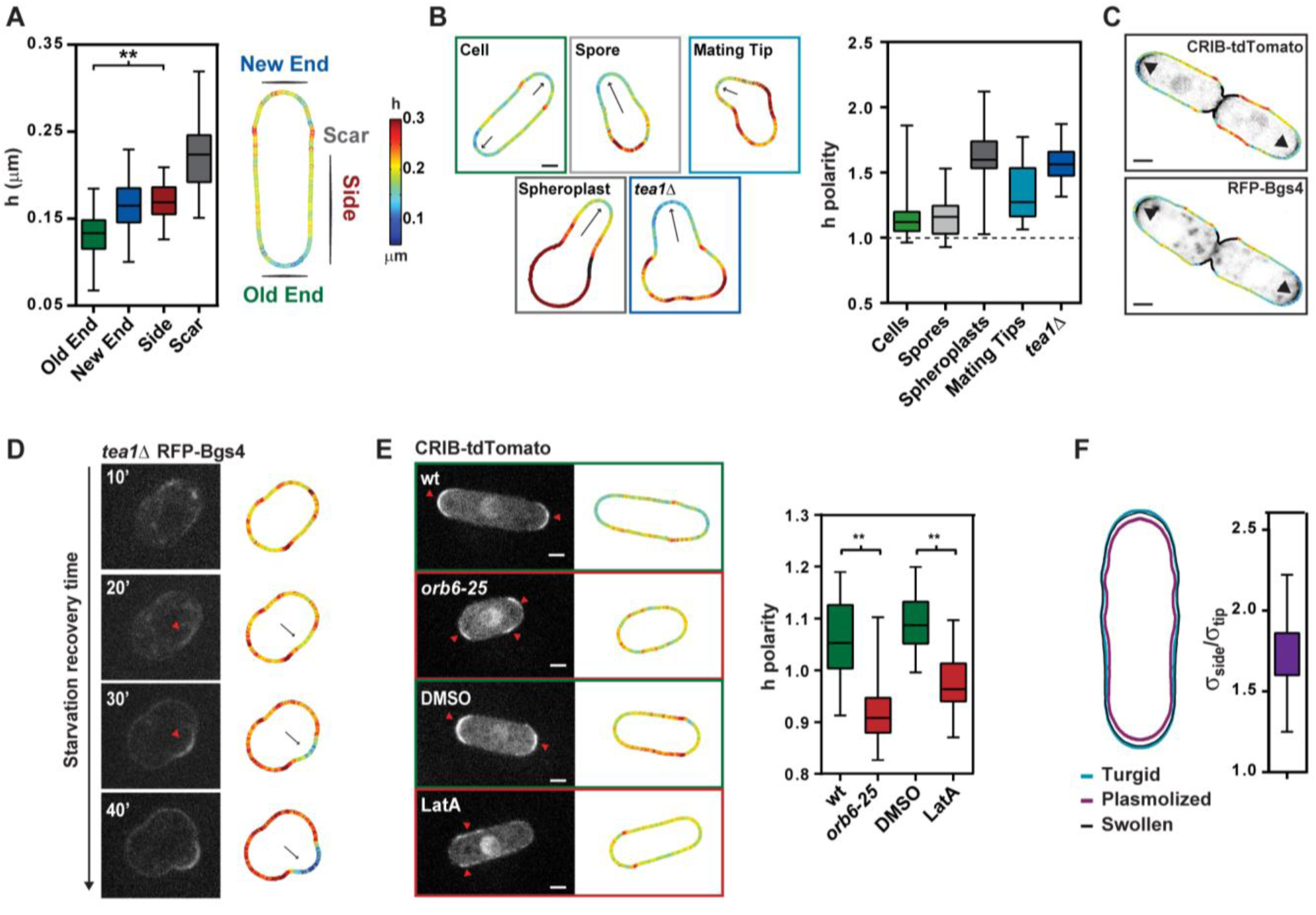
Cell polarity and cell wall thickness patterns. **(A)** Thickness color map of a typical interphase cell, and thickness values at the old end (n=50), new end (n=41), cell sides (n=51) and birth scars (n=47). **(B)** Representative thickness color maps and thickness polarity in wt interphase cells (n=41), outgrowing spores (n=24), mating tips (n=18), recovering spheroplasts (n=22), and *tea1Δ* cells recovering from starvation (n=29). Arrows indicate growth directions. h polarity is defined as a tip/side ratio in cells using an average of both tips values, and tip/back ratio in the other cases. **(C)** Superimposition of h map and polar growth markers (arrowheads). **(D)** Mid-slice confocal time-lapse and h maps of a *tea1Δ* cell recovering from starvation. Arrows indicate growth direction and arrowheads point at RFP-Bgs4 polar cap. **(E)** (Left) Midslice confocal image and h maps in the indicated conditions. Red arrowheads point at CRIB-td-Tomato polar caps. (Right) h polarity in wt (*n=30*) and *orb6-25* (*n=29*) grown 2h at restrictive temperature and in cells treated 30’ with DMSO (*n=21*) or 100 µM LatA (*n=26*). **(F)** (Left) Symmetrized cell boundary in turgid, plasmolyzed and computationally swollen state, used to compute local elastic moduli, Y, at cell tips and sides. (Right) Ratio of surface modulus, σ = hY between tips and sides (n=21 cells). Scale bars, 2 µm; Whiskers plots show median and full data set range. ** P < 0.0001. Black points in thickness maps correspond to positions in which thickness cannot be measured accurately.

Not surprisingly, these thin wall regions at sites of polar growth co-localized with the downstream polarity regulator GTP-Cdc42 (visualized with a CRIB domain fused to td-Tomato) and with the β-Glucan synthase Bgs4 (Cortes et al., 2005; Tatebe et al., 2008) (Figure 2C). Furthermore, time-lapse of polarized regrowth in *tea1Δ* cells recovering from starvation evidenced the stabilization of a Bgs4 polar domain, followed by concomitant CW thinning and emergence of a new growing tip (Figure 2D and Movie S3). Importantly, full depolymerization of actin with 100µM Latrunculin A or the use of an *orb6-25* thermo-sensitive allele affecting Cdc42-based polarity, both yielded the detachment of GTP-Cdc42 polar domains from cell tips, and consequent alteration of CW thickness patterns within tens of minutes (Bendezu and Martin, 2011; Das et al., 2009) (Figure 2E).

We also found that the CW was softer at growing cell tips. This was evidenced by computing CW elastic moduli, Y, which characterizes the bulk mechanical properties of the CW. To this aim, we plasmolyzed cells by rapidly adding 1.5M sorbitol to the medium in microfluidic flow channels which causes a rapid drop in turgor pressure, and consequent cell shrinkage (Abenza et al., 2015; Atilgan et al., 2015; Bonazzi et al., 2014). Using the changes in cell shapes between inflated and plasmolyzed states and local thickness values, we could extract values of CW elastic moduli at cell tips and at cell sides, to be Y_tip_=45.3 +/- 0.7 MPa; and Y_side_=72.5 +/- 1.15 MPa, yielding a ratio in surface moduli (σ=hY) of σ ^tip^/ σ s_ide_= 1,7+/-0.2 (Figure 2F and Figure S3C-S3D). Together, those results suggest that internal polarity which directs growth and CW synthesis causes sites of polar growth to have thinner and softer CW.

### Contribution of synthesis and growth to CW thickness at cell tips

Those findings prompted us to assess how CW thickness and assembly may be determined and regulated at sites of polar growth. We first considered a minimal theoretical model for thickness regulation at cell tips (Figure 3A). We posited that the CW thickens through synthesis and thins through cell elongation, which, given mass conservation yields a dynamic evolution of h_tip_:

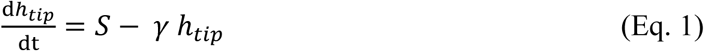

where S is a rate of wall synthesis per unit surface and γ the strain rate of the CW, which is related to the cell elongation rate, Gr, by the curvature radius at the tip, R_c_:

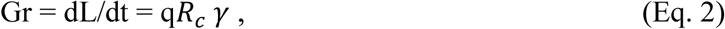

with L the length of the cell and q a numerical geometrical pre-factor.

**Figure 3.**
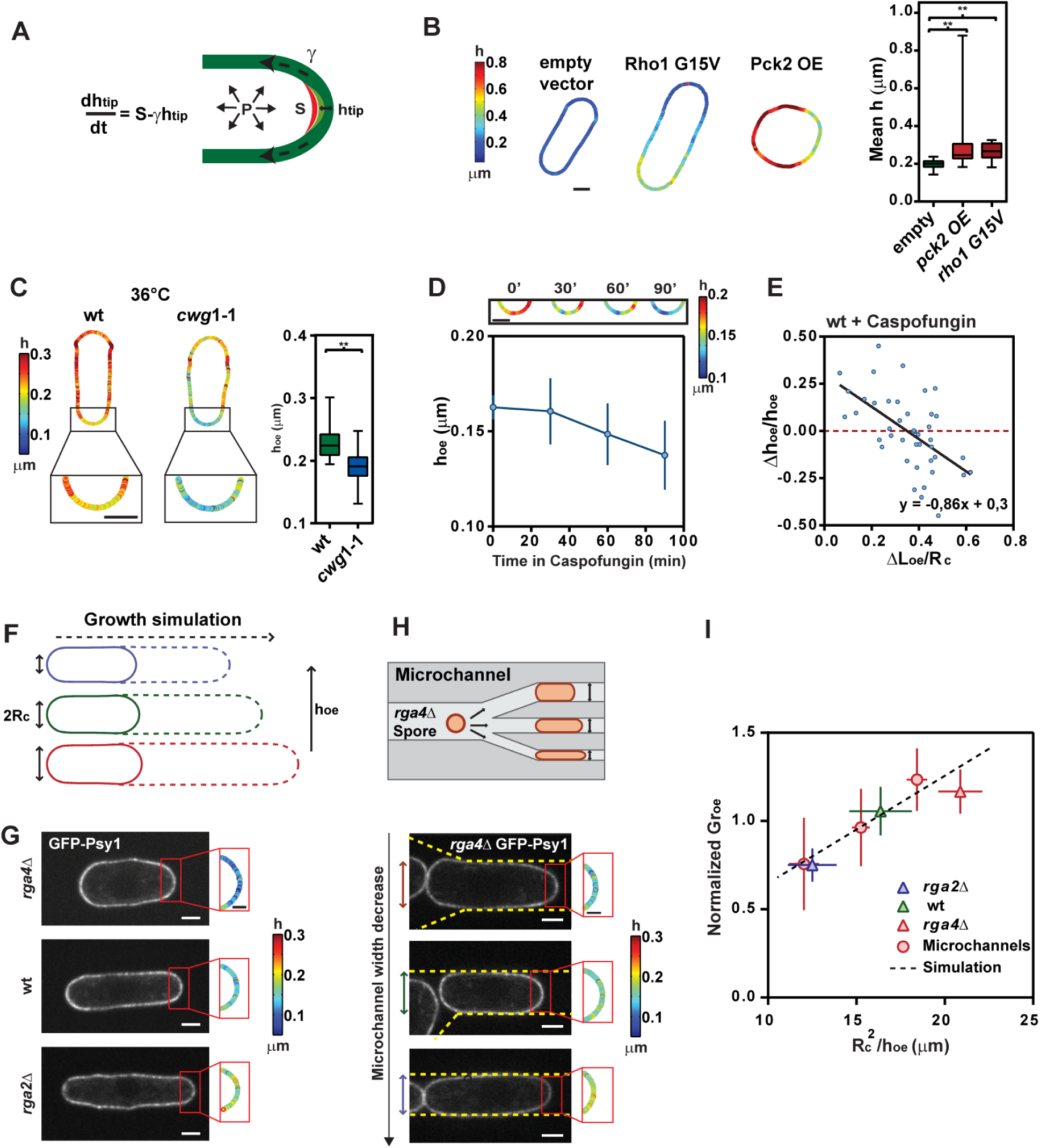
Dynamics of cell wall assembly at polar growth sites. **(A)** Simple model for tip thickness, h_tip_, dynamics. S is a rate of wall synthesis per unit surface, γ the strain rate and P the turgor pressure. **(B)** Representative color maps and mean h measurement of controls (n=59 cells), the constitutively active Rho1-G15V (n=27), and cells over-expressing Pck2 (n=59). **(C)** Local tip thickness in wt (n=27) and *cwg1-1* (n=29) grown at restrictive temperature. **(D)** (Top) Representative h_oe_ maps following Caspofungin treatment, and evolution of h_oe_ in time (Bottom, n=11 cells). **(E)** Δh^oe^/h_oe_, measured in Caspofungin treated wt cells and plotted as a function of ΔL/R^c^(n=44 in 12 cells). The black line is a linear fit. **(F)** Growth simulations: plain and dashed lines mark cell shapes at the beginning and the end of the simulation. **(G)** Midslice confocal images of typical *rga4Δ*, wt and *rga2Δ* cells expressing the membrane marker GFP-Psy1 and close-up local h color maps at the old end. **(H)** Schematic for diameter manipulation in microchannels, and midslice confocal images of *rga4Δ* cells grown in microchannels with different diameters and close-up local h color maps at the old end. The yellow dashed lines mark the border of the microchannels. **(I)** Normalized growth rate at the old end plotted as a function of R_c_^2^/h_oe_, for wt (n=36), *rga2Δ* (*n=*26), *rga4Δ* (*n=*37) and, *rga4Δ* cells grown in microchannels (*n=*33, binned on 11, 10, 13 cells). The dotted line is a fit of the growth simulation results. Whiskers plots show median and full data set range. Error bars correspond to SD. ** P < 0.0001. Scale bars 2 µm in confocal images and full cell color maps, and 1 µm in insets.

To assay the contribution of synthesis, we performed experiments to manipulate synthase activities. In cells overexpressing Pck2, a positive regulator of glucan synthesis, one major component of the fission yeast CW, we could detect significant thickening, up to ~900 nm in some cells (Arellano et al., 1999). Similar thickening was also observed by over-expressing a constitutively active Rho1 allele, *rho1-G15V* (Arellano et al., 1996) (Figure 3B). Conversely, a *cwg1-1* thermo-sensitive allele of the glucan synthase Bgs4, exhibited significant thinning at growing cell tips, consistent with previous TEM observations (Munoz et al., 2013) (Figure 3C). As a more direct assay, we also reduced CW synthesis with low doses of Caspofungin, a drug that specifically impairs β-glucan synthase activity (Martins et al., 2011). This treatment led to a net thickness decrease at growing ends over tens of minutes, eventually causing aberrant cell bulging at longer times (Figure 3D and movie S4).

Using those Caspofungin treated cells, in which synthesis is partially impaired, we also computed the contribution of growth to thickness variation. We measured the relative changes in thickness, Δh^oe^/h_oe_, which, given Eqs. 1 and 2, is predicted to linearly scale with -ΔL^oe^/R_c_. This analysis yielded a negative linear correlation with a slope of -0.81 (Correlation coefficient, r=-0.57), and a positive Y-intersect, which could correspond to remaining CW α-glucan synthesis in the presence of this drug (Figure 3E). Together, those data support a simple model for CW thickness evolution positively regulated by synthesis and negatively by growth.

### Influence of cell wall thickness on cell growth rates

As the balance between turgor and CW mechanics has been proposed to set growth rates in walled cells (Bonazzi et al., 2014; Minc et al., 2009), we sought to investigate the influence of thickness on strain and growth rates. Using a simple model for wall rheology (Minc et al., 2009; Rojas et al., 2011), we posited that the strain rate could be written as:

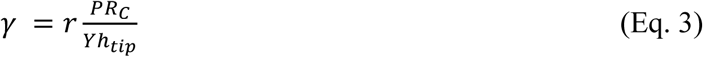

With r a wall remodeling rate, Y the elastic modulus of the wall, and P the turgor pressure (We here neglect a possible threshold in turgor for growth to occur). Given Eq. 2, this assumption implies that the elongation rate Gr scales with *Rc*^2^/*h*_*tip*_. This scaling was confirmed using 3D numerical simulations of a mechanical model of fission yeast growth (Abenza et al., 2015) (Figure 3F).

Experimentally, we used time-lapse movies to compute and plot elongation rates, as a function of initial tip curvature radii and CW thickness. To vary R_c_, we used wild-type cells and mutants with larger (*rga4Δ)* and smaller diameters (*rga2Δ*), as well as microfabricated channels to physically diminish the diameter of *rga4Δ* cells (Das et al., 2007; Villar-Tajadura et al., 2008; Zegman et al., 2015). This analysis yielded, as predicted, a robust linear dependency between Gr and *Rc*^2^/*h*_tip_ indicating that CW thickness could serve as a major determinant influencing elongation rates (Figure 3G-3I).

### Homeostasis in Cell wall thickness through strain mechanosensing

The above findings suggest that without regulatory layers, growth and thickness may negatively regulate each other, which could potentially lead to catastrophic situations of cell lysis through CW thinning or growth arrest through thickening. We thus computed tip thickness evolution in single cells over 1h, with a high temporal resolution of 6min, focusing on the old end. Strikingly, this revealed a thickness oscillatory behavior with phases of CW thickening, followed by thinning phases, and so on. Thickness on cell sides, was in contrast almost completely stable (Figure 4A-4B and Figure S4A-S4D). Those oscillating variations at cell tips, suggested the presence of a delayed feedback dynamically regulating CW thickness and assembly at growing tips.

**Figure 4.**
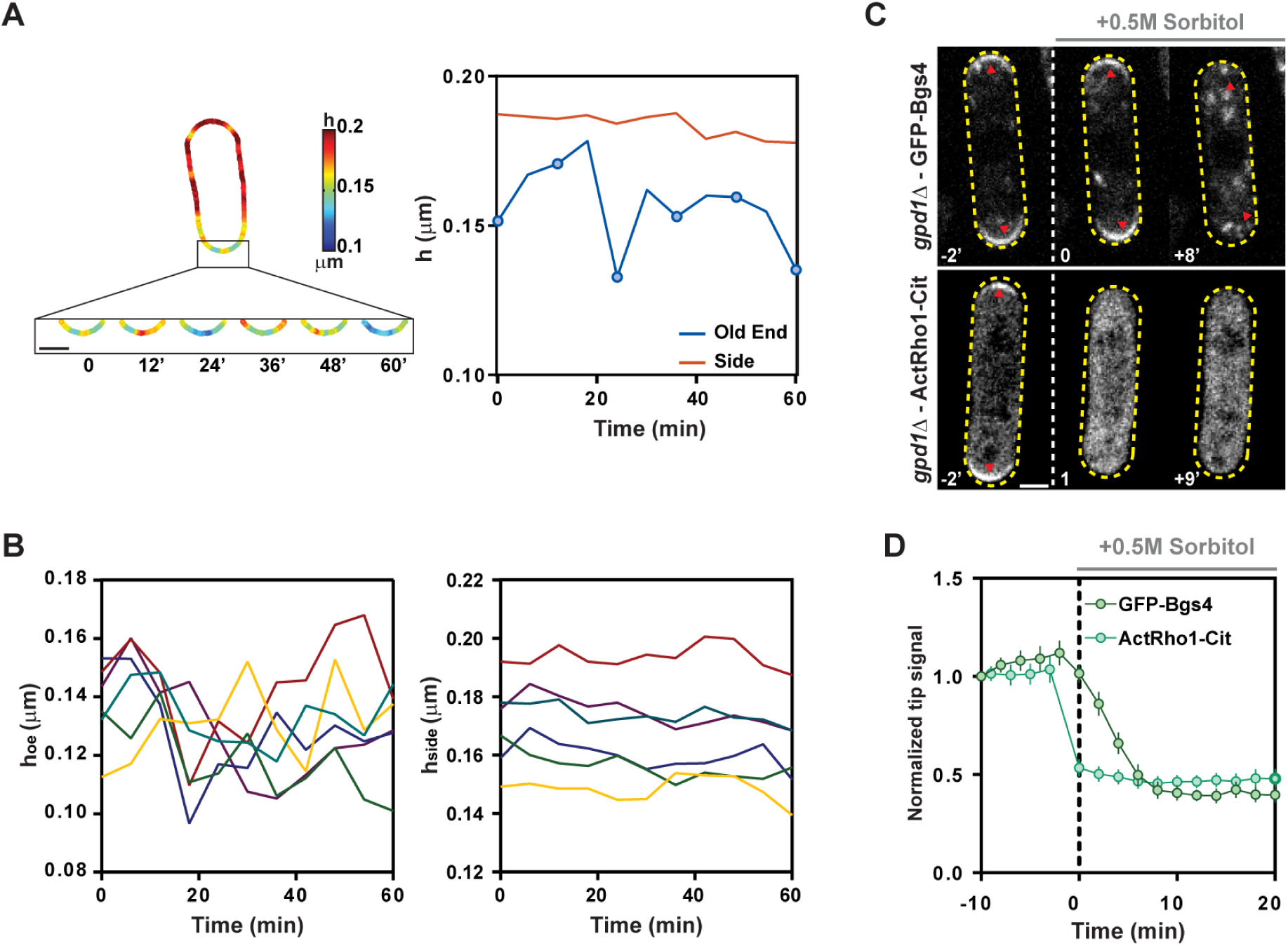
Cell wall thickness oscillations and strain rate-dependent CW synthase activation and localization. **(A)** Close up views on thickness maps at the old end of a growing wt cell, and corresponding plots of thickness at the old end and on the cell sides. Blue dots correspond to time points shown in color plots. **(B)** Evolution of thickness at the old end h_oe_ and on cell sides h_side_ imaged at 6 min interval in 6 wt cells (colors of the lines correspond to the same cell in both plots). **(C)** GFP-Bgs4 and Active-Rho1-Citrin polar domains (arrowheads) detachment in *gpd1Δ* cells following turgor reduction by addition of 0.5M sorbitol addition. **(D)** Evolution of normalized tip signal of GFP-Bgs4 (n=14) and Active-Rho1-Citrine (n= 18) in cells rinsed with sorbitol at t=0. Scale bars, 2 µm; Error bars represent SD. Scale bars 2 µm.

Based on reported observations in fission yeast and other cell types (Bonazzi et al., 2014; Nakayama et al., 2012), we assayed a potential positive feedback between growth, or equivalently strain rate, on polarized synthesis. To reduce strain rate, we rapidly decreased turgor, by rinsing cells with medium containing 0.5M sorbitol using a *gpd1Δ* background to prevent turgor adaptation (Minc et al., 2009). This caused an immediate growth arrest, and the rapid detachment of active-Rho1 polar domains (Davidson et al., 2015) followed by the detachment of Bgs4 domain over the subsequent minutes (Figure 4C-4D and Figure S4E). Those data suggest that CW synthase activity and localization may be actively sensitive to growth or strain rates.

To incorporate mechanical sensing in our model, we used a general delayed dependence, between strain rate and synthesis, of the form: S(t)=S_0_+λ*γ(t-T), with S_0_ a basal synthesis rate, λ a positive parameter characterizing the strength of the sensing and T a delay in the response between strain and synthesis, so that:

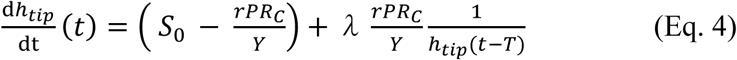

Using the evolution of old end thickness in single cells, we thus evaluated experimentally the dependence of the thickness rate, dh_oe_/dt (t_0_+T) computed at different delays, on the inverse of initial thickness 1/h_oe_ (t_0_), measured at various t_0_ in time-lapses. We found a positive correlation at delays starting around 12 min (r=0.4) which improved for longer delays, yielding highest correlations at delays around 30-40 min (r=0.8) (Figure 5A and Figure S5A-S5E). In this plot, the X-intercept corresponded to a homeostatic thickness of h_oe_* = 119 nm, close to the mean old end thickness at the population level. Importantly, similar homeostasic behavior was observed in *Cdc10-M17* and *Cdc25-22* cell cycle mutants respectively blocked in G1 or in G2 phase, ruling out putative contribution of cell-cycle transitions to this behavior (Figure 5B and Figure S5F-S5H). Solving the delayed differential Eq. 4 for h(t) yielded, in a range of parameter values, oscillatory behavior, reminiscent of experimental observations (Figure S5I-S5K and supplementary methods). Together those findings support the existence of a homeostatic mechanism, in which an overshoot (or an undershoot) in thickness is corrected by thinning (respectively thickening) the wall over the subsequent tens of minutes to maintain it around a narrow range of values.

**Figure 5.**
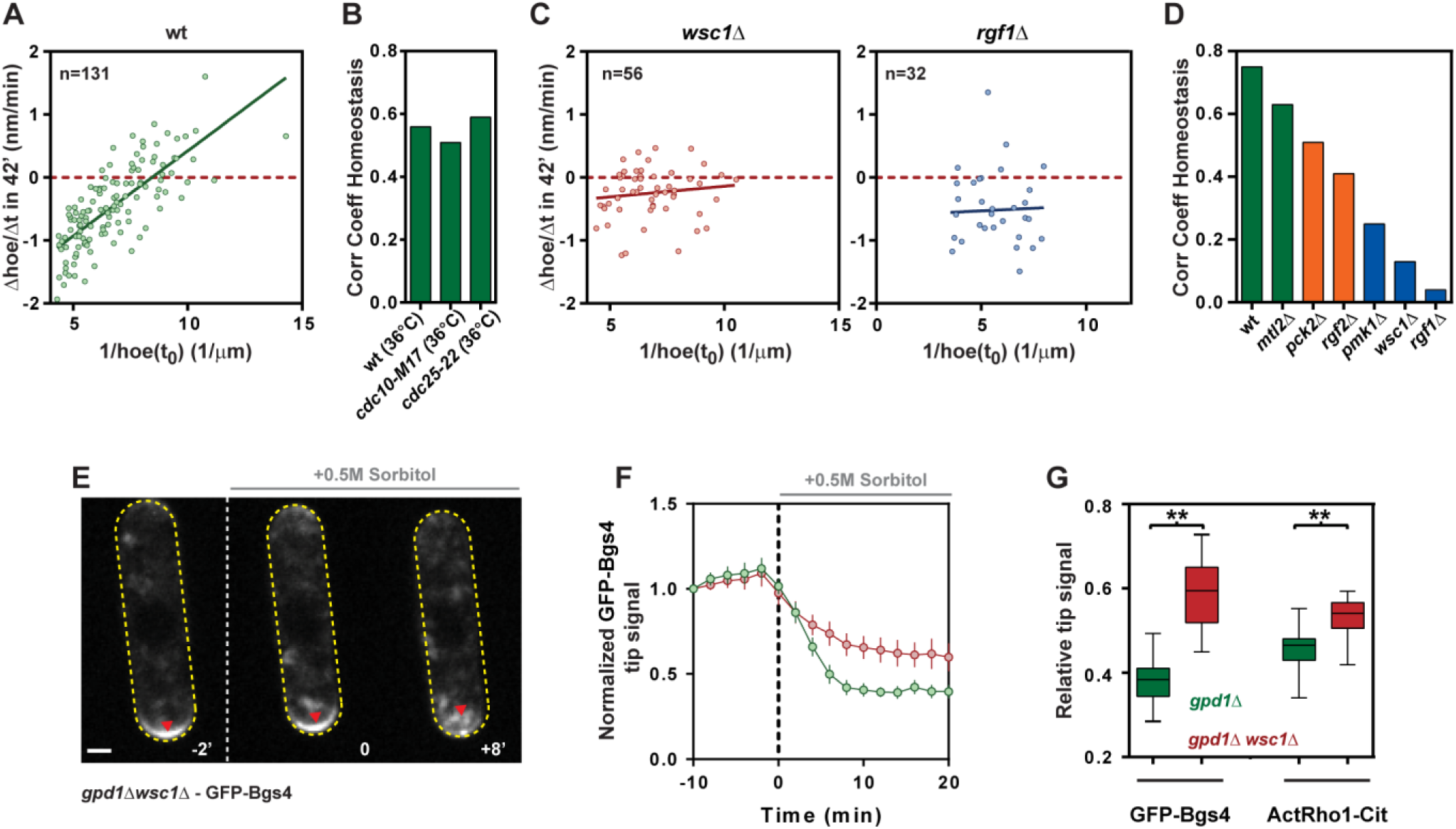
The Cell Wall Integrity pathway mediates cell wall thickness homeostasis. **(A)** Thickness homeostasis plot. Δh^oe^/Δt computed as (h^oe^(t_0_+42’)-h_oe_(t_0_))/42, for various t_0_ and plotted as a function of 1/ h_oe_(t_0_), for wt (n=131 cells in 25 cells). The plain line is a linear fit. **(B)** Correlation coefficient for thickness homeostasis for cells of wt (n=82 in 21 cells), *cdc10-M17* (n=48 in 13 cells) and *cdc25-22* (n=80 in 20 cells), grown at restrictive temperature (36°C). **(C)** Thickness homeostasis plot for *wsc1Δ* (n=56 in 14 cells) and *rgf1Δ* (n=32 in 8 cells) mutants. Palin lines are linear fits. **(D)** Correlation coefficient for thickness homeostasis for wt (n=59 cells), *mtl2Δ* (n=56 in 14 cells), *pck2Δ* (n=40 in 10 cells), *rgf2Δ* (n=52 in 13 cells) and *pmk1Δ* (n=39 in 10 cells). **(E)** Defect in GFP-Bgs4 detachment in response to turgor reduction in a *gpd1Δwsc1Δ* double mutant. Note the remaining signal as compared to a *gpd1Δ* single mutant shown in Figure 4C. **(F)** Evolution of normalized tip signal of GFP-Bgs4 (n=19) in *gpd1Δ* and *gpd1Δwsc1Δ* cells rinsed with sorbitol at t=0. **(G)** Ratio between tip signals after and before sorbitol treatment in *gpd1Δ* and *gpd1Δwsc1Δ* cells (n=14, 19, 18, 18). Whiskers plots show median and full data set range. Scale bars, 2 µm; Error bars represent SD.

### The Cell Wall Integrity (CWI) pathway mediates strain mechanosensing for thickness homoestasis

A prime candidate system which may sense CW properties and influence synthesis is the cell wall integrity pathway (CWI), a signaling cascade involved in general CW stress response (Perez and Cansado, 2010). In fission yeast, this pathway is thought to be activated at the cell surface, by the putative trans-membrane stress sensors Wsc1 and Mtl2 which interact with the CW plausibly through long serine/threonine rich domains extending in the CW matrix (Cruz et al., 2013). Those factors signal downstream to Rho-GEFs Rgf1 and Rgf2 which may directly regulate the Rho-GTPases Rho1 and Rho2 for glucan synthase activation (Garcia et al., 2009a; Garcia et al., 2006; Mutoh et al., 2005). The CWI also triggers the activation of gene transcription for CW repair, through the Pmk1 MAPK cascade (Garcia et al., 2009b; Perez and Cansado, 2010). By performing a candidate screen on single mutants of the CWI, we found that *wsc1Δ, rgf1Δ* and *pmk1Δ*, were severely defective in thickness homeostasis, with changes in thickness being mostly independent on previous thickness values (Figure 5C-5D and Figure S6A-B). In addition, the detachment of Bgs4 and active-Rho1 polar domains as a response to pressure and strain reduction in a *gpd1Δ wsc1Δ* mutant was much less pronounced than in a *gpd1Δ* alone, suggesting that Wsc1 may directly probe CW strain rate to tune synthesis through the CWI (Figure 5E-5G and Figure S6C).

We then assayed if the implication of the CWI pathway in thickness homeostasis could account for the lysis defect of some CWI mutants during normal growth (Cruz et al., 2013; Garcia et al., 2006). By imaging thickness evolution in *rgf1Δ*, we observed in ~30% of cases, cells that started with a relative thick wall and little growth, which switched incidentally to a phase of unusually rapid growth concomitant with rapid CW thinning eventually yielding to tip lysis and cell death (Figure 6A-6B). We noted, however, that the thickness at which lysis occurred in some of those cells could have values similar to wild-type tips. As the failure strain of the CW yielding to lysis may depend on the surface modulus, σ=Yh, which has contributions from both thickness and elastic modulus, we suspect that those cells have lower bulk modulus, arising from mis-regulated CW composition. This was supported by the similar growth rate values reached before lysis, which is predicted to inversely depend on the surface modulus (Figure 6B). Similar lysis phenotype was also observed in *wsc1Δ*, but to a much lesser extent which precluded a careful analysis of thickness dynamics. Together those findings demonstrate that the CWI may directly probe cell wall strain rates to adjust synthesis, thereby coordinating growth and CW assembly needed for cell survival (Figure 6C).

**Figure 6.**
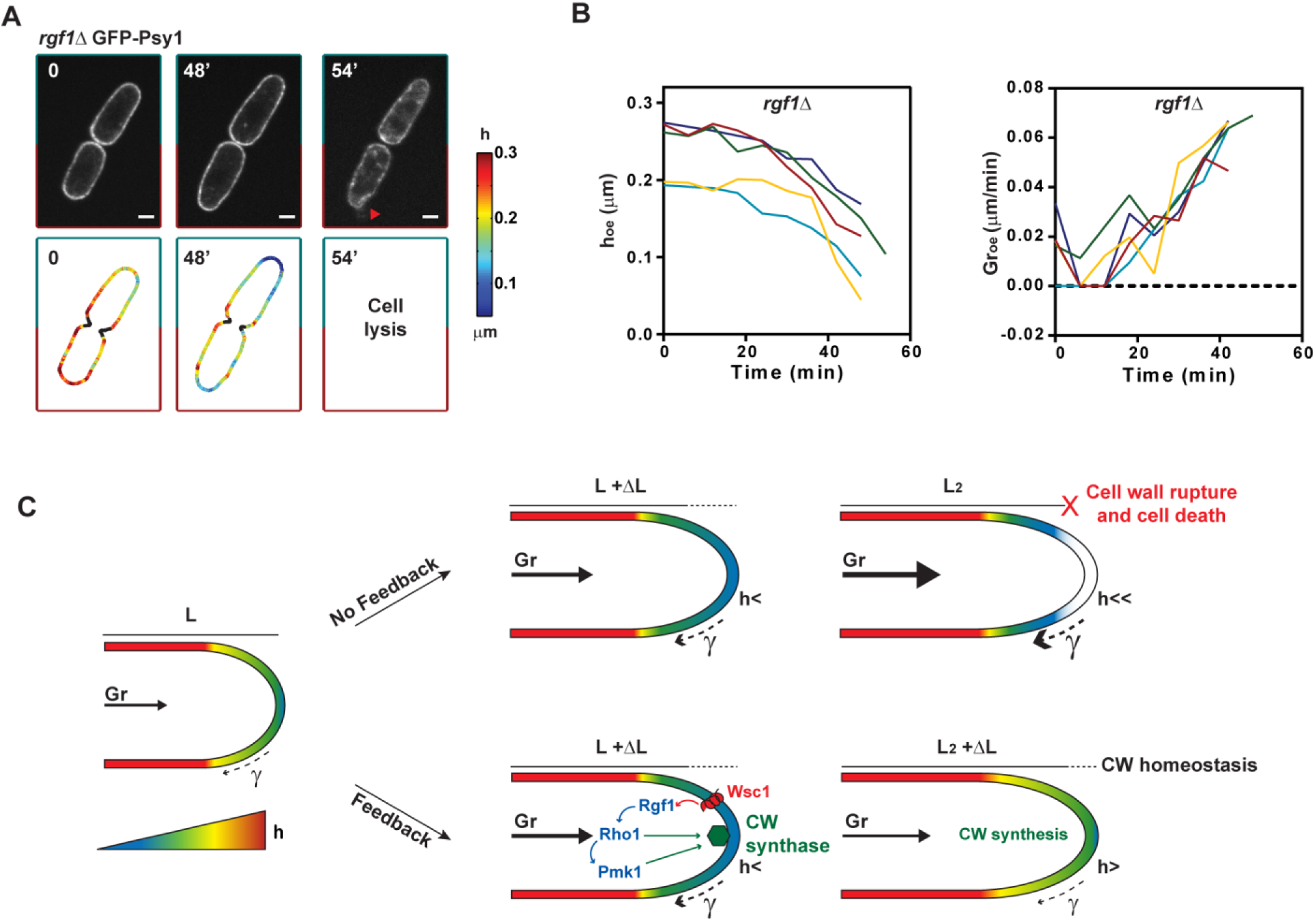
Implication of thickness homeostasis for cell survival during normal growth. **A** Time-lapse of GFP-Psy1 and thickness maps for a representative lysing *rgf1Δ* cell. The arrowhead points at membrane leaking at cell tips after lysis. Evolution of old end thickness h_oe_ and growth rates Gr_oe_ in lysing *rgf1Δ* cells. The last point is recorded before lysis. **(C)** Mechanosensing-based feedback for CW thickness homeostasis during cell growth. In absence of feedback, growth causes cell wall thinning which promote faster growth and further CW thinning and so on, until the CW ruptures and cells lyse. The cell wall integrity pathway probes strain rate (or equivalently growth rates) through the surface protein Wsc1 that promotes the activation (directly or indirectly) of the Rho1-GEF Rgf1 to activate of Rho1 and cell wall synthase (CWs). Rho1 can directly act on synthesis or through the Pmk1 MAPK cascade. As a result, thickness increases, thereby reducing strain rate, yielding to a dynamic homeostatic system for CW mechanical properties and cell integrity. Scale bars, 2 µm; Error bars represent SD. **P < 0.0001

## DISCUSSION

The mechanical properties of the CW and turgor values underlie walled cells growth and shapes, and have been redundantly used as tunable cellular properties to survive, colonize, infect or reproduce (Bastmeyer et al., 2002; Dagdas et al., 2012; Dudin et al., 2015; Harold, 2002; Keegstra, 2010; Lew, 2011; Silhavy et al., 2010). This importance contrasts with the limited knowledge we have on the dynamic regulation of those mechanical parameters in growing and dividing cells. By developing and validating here the very first approach to monitor CW thickness in live cells, we evidence key contributions of the polarized dynamics of the CW for growth and viability. Our approach overcomes numerous limitations of EM studies, such as alterations in cell shape and CW caused by fixation (Osumi, 2012), and brings the possibility to visualize the CW in a planar section, in large populations of living cells. We foresee that this approach could serve as a novel standard for studying CW function in processes such as growth, reproduction or infection in bacteria, fungi or plants, with potential therapeutic values for anti-fungal and antibiotic chemical screens.

### Spatio-temporal Cell Wall dynamics

One important output of our study, is to find that CW thickness and mechanics are polarized, with growing poles being typically twice softer and thinner than the rest of the cell. Those mechanical anisotropies, which here derive from internal polarity; have long been speculated to be required for tip elongation and rod-shape morphogenesis (Boudaoud, 2003; Drake and Vavylonis, 2013; Rojas et al., 2011). Patterns of wall elasticity may also influence the morphogenesis of other tip growing cells (Ma et al., 2005; Yanagisawa et al., 2015), as well as that of multicellular plant tissues (Kierzkowski et al., 2012; Milani et al., 2014; Peaucelle et al., 2011). Given the large variety of shapes found in fungal species and mutants, it will be interesting to systematically compare spatial patterns of CW mechanics and thickness with cell shape parameters, to identify generic biomechanical principles guiding the morphogenesis of walled cells. Animal cells also exhibit anisotropic mechanical properties needed for shape changes, cell migration or tissue morphogenesis, which rest on the organization of internal polarity and cytoskeleton (Levayer and Lecuit, 2012; Mogilner and Keren, 2009). Thus, our data reinforces a concept that morphogenesis may primarily emerge from spatial surface mechanical properties patterned from intracellular biochemical organization.

We also discovered that the CW is highly dynamic at sites of polar growth, with variations in thickness amplitude reaching up to 30% of the mean, on a time-scale of few minutes. Thickness on cell sides appeared comparatively more stable, although we did note some patterns of thickness translating away from cell tip (movie S1 and S2), plausibly reflecting CW material flowing during cell growth (Abenza et al., 2015). We suggest that thickness oscillations are indicative of a homeostatic system correcting thickness changes overtime, needed to maintain CW mechanical parameters in an optimal range for growth and viability. Although the current state of the art precludes from dynamically computing changes in elastic moduli in growing cells, we suspect that similar homeostatic systems could also influence composition and bulk elasticity in addition to thickness. CW mechanical oscillations have been speculated to exist in rapidly growing cells such as pollen tubes and fungi; and to be related to cycles of Rho-GTPase activation or Calcium signaling and variations in growth rates (Qin and Yang, 2011; Rojas et al., 2011). Mechanical oscillations may thus represent a general read out for negative-feedbacks acting at the biochemical or biomechanical level, instrumental to confer robustness to growth and viability (Howell et al., 2012).

### Mechanical homeostasis of the CW by the CWI

By screening through mutants, we uncovered a primary role for components of the CWI in probing CW strain and cell growth. Although the role of the CWI in general CW stress response is well established in yeast, fungi and plants, to our knowledge, those are the very first direct evidence of a direct role in dynamically controlling CW properties during normal cell growth. The implication of Wsc1 in thickness homeostasis and in probing CW mechanical strain to regulate synthase polar domains localization and activity, provides strong support that this protein may act as a direct CW mechanical sensor. Wsc1 localizes in patches at cell tips, and has a predicted serine/threonine rich domain spanning the CW (Cruz et al., 2013). It remains a puzzle to understand how such configuration could directly probe lateral strain in the CW. Studies of *S. Cerevisiae* orthologues have proposed that WSC clustering may underlie mechanosensory activities (Heinisch et al., 2010); further work will be needed to understand how those domains may probe the dynamic mechanical properties of the CW. Mutants in the second fission yeast stress sensor, Mtl2, did not exhibit strong defects in homeostasis, suggesting a dominant function of Wsc1, here. This may be in agreement with the more diffuse localization of Mtl2 all around the cell surface (Cruz et al., 2013). Surprisingly, we found that loss of Rgf2, which has been suggested to physically interact with Wsc1 (Cruz et al., 2013), only had a mild effect on wall thickness homeostasis. *Rgf1Δ* mutants exhibited the most pronounced effect, which may account for its severe lysis phenotype. This suggests a potential link between Wsc1 sensing and Rgf1 activation in wall homeostasis, and a dominant role for Rgf1p in activating Rho1 in this response (Garcia et al., 2009a). However, we note that 70% of *rgf1Δ* mutant cells still survived, suggesting that a secondary system, likely based on Rgf2 may maintain some form of mechanical homeostasis, plausibly at longer time-scales. Finally, we also found that Pmk1 was required for homeostasis. This suggest that proper homeostasis involves at least partially the expression of a set of regulatory genes for CW construction. This would imply that the Pmk1 cascade acts at short time-scales of several minutes, similar to turgor pressure-regulating Sty1/Hog1 MAPK (Miermont et al., 2011). Thus, the dynamic regulation of walled cells mechanical properties, through rapid adaptation systems, appear as a central element to support viability not only in response to stress, but also during normal growth and division.

### Growth control through mechanochemical systems

The notion that surface mechanics can alter the core spatio-temporal organization of cells, through mechanosensing systems is becoming increasingly important in biology (DuFort et al., 2011). Examples range from the regulation of cell polarity, migration, cell division up to tissue homeostasis (Bonazzi et al., 2014; Fink et al., 2011; Houk et al., 2012). In here, we uncovered a core function of mechanosensing in the control of growth rates. In that view, growth is directly perceived as a mechanical signal straining the CW, without any requirement for interactions with neighboring cells or cell environment. When a cell begins to elongate rapidly, the strain rate increases, which turns on synthase activation by the CWI to assemble more wall, and *vice e versa*. This homeostatic system thus compensates surface material dilution associated with growth, to maintain its thickness around values compatible with survival. As thickness influences elongation rates, this system may also work towards controlling growth rate values around a defined range. Those findings thus have important implications for cell cycle control, as well as size homeostasis, which are both influenced by growth rate values. We speculate that cells may have evolved built-in mechanical properties and sensing elements which allow them to control expansion speeds adapted to their life-styles. Future work studying similar problems of surface expansion control will inform on how those elements have been modified to guide the growth of rapidly expanding cells like fungi and plant, or slowly growing cells like those of animals.

## AUTHOR CONTRIBUTION

V.D, A.H., H.D.B, R.L.B and N.M. performed experiments. H.T., D.E. and N.M. developed image analysis scripts. E.C. and A.B. performed modelling. V.D., A.B. and N.M designed the research and wrote the manuscript.

## ACKNOWLEDGEMENTS

The authors acknowledge P. Perez, Y. Sanchez, H. Valdivieso, T. Toda, T. Nakamura (YGRC/NBRP Japan) and J.Q. Wu, for sharing material. We acknowledge the ImagoSeine facility, a member of France BioImaging (ANR-10-INSB-04). Our laboratory is supported by the CNRS and grants from the FP7 CIG program, ITN ‘‘FungiBrain’’, and the European Research Council (CoG Forcaster no. 647073).

## STAR METHODS

### EXPERIMENTAL MODEL AND SUBJECT DETAILS

#### Yeast strains, media and genetics

Standard methods for *Schizosaccharomyces pombe* media and genetic manipulations were used (http://www-bcf.usc.edu/~forsburg/). Strains used in this study are listed in Key resource table below. Cells were grown at 25°C in yeast extract plus 5 supplements (YE5S) media unless otherwise indicated. Over-expression driven from the plasmids, was induced by growing cells at 25°C in Edinburgh minimal medium (EMM) + supplements without thiamine for 24-36 h for Pck2p Over expression, and 48h for the membrane probe GFP-RITC and for ActRho1-Cit. Starvation of wild-type (wt) cells was induced by over-growing cells in liquid YE5S for an additional 16 h after they had reached stationary phase (OD_600_>1). Starvation of *tea1Δ* cells was induced by growing cells on YE5S plates for > 3 days, and subsequently transferring them in YE5S liquid media for 2-3h before observation or directly on 2% YE5S agar pads for time lapse. Temperature sensitive alleles, *cwg1-1* and *orb6-25* were grown in liquid culture at permissive temperature (25°C) and switched to restrictive temperature (36°C) for 6h and 2h, respectively, before observation. Spores were obtained from homothallic h90 strains, or from heterothallic crosses. Freshly growing cells were placed on malt extract (ME) solid media for 3 days. Mating mixtures were then digested for 1h at room temperature in a 1/200 glusulase solution in water to kill vegetative cells, washed three times in water, and incubated for >7h in YE5S before observation (Bonazzi et al., 2014). Spheroplasts were generated from exponentially growing cells in YE5S, which were washed twice in digestion buffer (0.1M Citric Acid 0.1 M NaCitrate pH = 5.8; 1.2 M Sorbitol) and subsequently treated with Lallzyme (0.1 g/ml) during 1h at room temperature. Newly formed spheroplasts, were washed twice in YE5S+1.2 M Sorbitol, and let to regenerate at 25°C for >16h, until outgrowing protrusions appeared (Flor-Parra et al., 2014). To image mating protrusions, homothallic cells growing exponentially in liquid YE5S were placed on solid ME until mating protrusion appeared (>5h). Cells were then harvested, labeled and imaged directly.

#### Pharmacological inhibitors

Cell wall digestion was achieved by incubating cells with 5000U/ml Zymolyase (ZymoResearch) at room temperature, and imaged immediately after treatment and 1h after. LatrunculinA (Sigma) was used at a final concentration of 100 µM from a 100X stock in DMSO. Cells were incubated for 30’ at room temperature before imaging. For Caspofungin experiments, cells were plated on 2% agar pads made with YE5S+10 mg/ml *Gs*-IB_4_-Alexafluor647 and a final concentration of 5µM Caspofungin.

### METHOD DETAILS

#### Microscopy

Live-cell imaging was performed on two different inverted spinning-disk confocal microscopes equipped with motorized stages, automatic focus and controlled with MetaMorph^®^ (Microscopy Automation & Image Analysis Software). The first one (Nikon Ti-Eclipse), is equipped with a Yokogawa CSU-X1FW spinning head, and an EM-CCD camera (Hamamatsu), a 100× oil-immersion objective (Nikon CFI Plan Apo DM 100×/1.4 NA) and a 2.5× magnifying lens, yielding a pixel size of 43 nm. The second one (Leica DMI8), is equipped with a Yokogawa CSU-W1 spinning head, and a sCMOS Orca-Flash 4 V2+ (Hamamatsu) a 100× oil-immersion objective (Leica Plan Apo DM 100×oil/1.4 NA), yielding a pixel size of 70nm.

Image registration was performed by imaging a slide containing a solution of 0.2 µm TetraSpeck™ microspheres (Thermofisher). A field containing one bead, was moved sequentially using the Metamorph function “ScanStage” coupled with a home built plugin to allow multidimensional acquisition. This generated an array of fluorescent spots imaged at multiple wavelength, for image registration (Figure S1A, see below). Alternatively, single images of a dense field of non-aggregated beads, could be used to generate the vectorial map. For cell wall thickness measurements, cells were pre-labeled in growth media containing 5 mg/ml of labelled lectin from *Griffonia simplicifolia Gs*-IB_4_-Alexafluor647 or *Bs*-IB_4_-TRIC (Horiseberger and Rosset, 1977). For single-time imaging, cells were placed between a glass slide and a coverslip and imaged within 20 min. For time-lapse imaging and for the experiments, cells were placed on 2% agar-pads made of YE5S containing 10 mg/ml *Gs*-IB_4_-Alexafluor647 and covered with a coverslip. We detected small differences in thickness between experiments performed in glass slides (202±19 nm, n=169) and in agar pads (177±28 nm, n=66), with glass slides generally yielding thicker values, which could be attributed to small cell deformation or starvation. For this reason, all experiments done in mutants, drugs or by modifying protocols are associated with a control measured in the same manner. Comparison with electron microscopy was performed on agar pads. Imaging was generally performed at room temperature (22-26 °C), with controlled humidity (>30%). For temperature-sensitive mutants, the temperature of the sample was kept at 36 °C by using either a temperature controlled cage or an objective heater (Bioptechs).

#### Transmission Electron Microscopy

For high pressure freezing (Figure S3A), cells were grown in liquid and harvested on a vacuum filter. The pellets were transferred to 100µm deep flat carriers (Leica Microsystems). Samples were then cryo-immobilized by HPF (EM-Pact2, Leica microsystems). Freeze substitution was performed with an AFS2 (Leica Microsystems) in 2% Osmium, 0,1% Uranyl acetate in pure acetone following protocols described in (Murray, 2008). Samples were rinsed three times with acetone and infiltrated with gradually increasing concentrations of an Epon resin mix (Agar scientific) and polymerized for 24h at 60°C. Ultrathin (70 nm) sections were generated with an ultramicrotome (UC6, Leica Microsystems) and collected on formvar/carbon-coated grids. Sections were post-stained by aqueous 2% uranyl acetate and lead citrate before imaging.

For chemical fixation (Figure S3A), samples were fixed with 2% glutaraldehyde in Phosphate Buffer 0.2M buffer, for 1h at room temperature followed by overnight fixation at 4°C and post-fixed with 2% osmium tetroxide in water. Samples were dehydrated through a graded series of ethanol and propylene oxide and embedded in epoxy resin. Ultrathin (70 nm) sections were generated with an ultramicrotome (UC6, Leica Microsystems) and collected on formvar/carbon-coated copper grids. Sections were then post-stained by aqueous 4% uranyl acetate and lead citrate.

For Correlative Light Electron Microscopy (CLEM) cells were immobilized on specialized bottom glass petri-dish with micro-grids for image position registrations (MatTek) (Figure S3D). Dishes were pre-treated with 1mg/ml poly-lysine and 0.1 mg/ml *Bs*-IB_4_-lectine and rinsed with water and YE5S. Cells, pre-labeled with *Gs*-IB_4_-Alexafluor647, were placed on the dishes, let to sediment and stick for ~15 min, and were subsequently rinsed vigorously with YE5S to detach non-sticking cells. Cells were then fixed with 2% glutaraldehyde in YE5S media for 2h at 4°C. Cells were then imaged in light microscopy and their positions were recorded using marks on the micro-grid. The samples were post-fixed with 2% osmium tetroxide in water and dehydrated through a graded series of ethanol and embedded in epoxy resin, which tends to stick to the micro-grid. Ultrathin (70 nm) sections of selected regions of the micro-grid, were generated with an ultramicrotome (Ultracut UC6, Leica) and collected on formvar/carbon-coated copper grids. Sections were then post-stained by aqueous 4% uranyl acetate and lead citrate. We observed often a partial cell deformation recorded in TEM, plausibly due to resin embedding. In spite of this the cell wall remained intact in most cases. All samples were observed in a Tecnai12 (FEI, The Netherlands) transmission electron microscope at 80 kV equipped with a 1K×1K Keen View camera.

#### Microfabricated channels

Microchannels used to manipulate cell diameter, were fabricated in PDMS from a silicon wafer printed with classical soft-lithography procedures, as described in (*24*) (Figure 3H). Homothallic *rga4Δ* spores were inserted into the PDMS microchannels, let to germinate for >20h at 25°C, and subsequently rinsed with YE5S + *Gs*-IB_4_-Alexafluor647 for 2h before imaging.

#### Turgor pressure manipulation by sorbitol addition

For sorbitol treatments cells were pre-labeled in growth media containing 5 mg/ml of labelled lectin *Gs*-IB_4_-Alexafluor647, then placed in PDMS microfluidic chambers between a dialysis membrane and a coverslip, which allowed live fluid exchange (Charvin et al., 2008). Cells were left in the chamber 40 minutes for adaptation before starting the assay. For Young modulus estimations cells were imaged, then immediately rinsed in the same media containing 1.5 M sorbitol and imaged again within the next 4 minutes. For growth rate manipulation, cells were imaged every 2 minutes for 10 minutes in YE5S + *Gs*-IB_4_-Alexafluor647, then washed in the same media containing 0.5M Sorbitol and imaged for 30-60 minutes.

### QUANTIFICATION AND STATISTICAL ANALYSIS

#### Image analysis

All image analysis scripts developed in the context of this study, can be made available upon request.

##### Cell Wall thickness measurement

The analysis pipeline to compute CW thickness used was fully automated with a manual check option for each step. All the analyses were performed using custom scripts written in MATLAB (Mathworks, R2013a equipped with Image Processing and Statistics toolboxes). Initial cell segmentation was performed using either fluorescent membrane or lectin signals. The signal was first binarized using a threshold determined with the graythresh function. The cell contour was defined as the most outer pixels of the binarized signal. The coordinates corresponding to the points along the cell contour were then equally spaced with an interval of 1 pixel using a spline function. Typically, a single cell contour was represented by ~ 600-800 points for a pixel size of 43 nm. At each point of the contour, a 40 pixels long line perpendicular to the cell contour was defined as the normal to the tangent of the curve determined using 11 neighboring points (Figure 1A first panel, blue line). The membrane and lectin signals were scanned along these lines, leading to one dimensional intensity profiles for each signal at each position around the cell contour (Figure 1A second panel). The center of each intensity profile was identified by fitting a Gaussian function on 11 points around the point with maximum intensity. Each of these centers defined the XY coordinates of membrane and lectin along the contour of the cell. To correct for chromatic shifts, the coordinates of the wall were changed using the position-dependent chromatic shift vectorial map (Figure S1A, see next section). This first set of analyses defines the precise in-plane coordinates of both outlines of cell membrane and lectin signals (Figure 1A, third panel). Subsequently, novel lines perpendicular to the lectin boundary spaced by 1pixel and were defined, following the same procedure as above (In Figure 1A second panel, only 1 out 10 of those lines are drawn, for clarity). The position of the cell membrane along the normal lines was defined as the intersection point between the normal line and the cell membrane. Finally, the local cell wall thickness h was computed as the distance between lectin and membrane coordinates along the normal line (Figure 1A third panel insert). Lateral resolution was estimated based on the lateral precision with which we could detect birth scars (~500nm) (Figure 1A fourth panel, Figure 1C and Figure S2E-S2G).

##### Chromatic shift correction

To correct for the chromatic positional XY shifts between the channels used for the lectin and membrane signals across the field of view we used a “point-by-point correction” approach. In contrast to standard “whole image” alignment approaches (Clark et al., 2013), this produces a continuous vector field containing shifts for all pixels in the field of view, and can thus account for non-uniformity in the positional drift. Starting from arrays of 200nm beads imaged at different wavelengths (see above), we first detect the particles positions with a sub-pixel precision (~20-25 nm) using a modified Matlab script available from M. Kilfoil lab (http://people.umass.edu/kilfoil/). The positions of the particles in the two channels (X1, Y1) and (X2, Y2) are then used to compute a sparse map of point-by-point shift vectors, defined as: V = (Vx, Vy) = (X2 – X1, Y2 – Y1) (with one channel used as the reference and features in the second one are being aligned) (Figure S1A right panel). This sparse array is then used to generate a continuous point-by-point shift map. For this, each of the two 3-dimentional sets of data: (X1, Y1, Vx) and (X1, Y1, Vy) are fitted with a plane, which allows for computing a point-to-point shift vector Vi at any position (Xi, Yi) with sub-pixel resolution close to the particle center detection limit (~20-25nm) (Figure S1A). This map is finally used as an input in the cell wall thickness measurement scripts described above (Figure S1C).

To assess the chromatic shifts in Z, we imaged a 3 µm deep Z stack with 10 nm spacing between slices of the same 200nm beads in 5 different positions in the field of view in two wavelengths channels. To reproduce the distance of the cell center from the cover glass (about 2 µm), beads were embedded in 2% agar pad, and only beads at about 2 µm from the glass surface were imaged. For each stack the fluorescent peaks were detected using a Gaussian fit, and the shift computed from the distance between the peaks. We found that this shift ranged typically from 0.1 to 0.4 µm, which corresponds to a projected shift in thickness smaller than 30nm that we neglected for the measurement (Figure S1B).

##### Generation and analysis of simulated images

Simulated images were generated as described in (Clark et al., 2013), using custom scripts written in MATLAB. The simulation starts with two dimensional data, corresponding to a simple spherical cell shape of 3 µm diameter, drawn with a 5 nm resolution. The simulated cell has a cell membrane thickness of 15 nm and a cell wall thickness of 200 nm. To best represent the experimental imaging conditions, the cell membrane was assumed to be uniformly stained, and only 5 nm of the most outer part of cell wall was uniformly stained in another color. To account for the experimental noise and finite background signals inside and outside of the cell, two different Gaussian noises, were added. The mean and standard deviation of the noise was determined so that the noise in the final image matches typical experimental signals. The original image was then convoluted with a Gaussian function with a standard deviation of 200 nm corresponding to the point spread function. The smoothed data were sampled at 40 nm to generate the final pixelized image. To systematically test for the effects of finite signals in the cytoplasm, we kept the standard deviation of the noise and changed only its mean values. The generated images were analyzed using the same scripts as for the experimental data described above.

The simulation results suggest that the inside-outside difference in the background signal could potentially influence cell wall thickness measurement (Figure S1D). By running a systematic modification of background difference between inside and outside, and signal/noise ratio in simulations, we thus computed the influence of those parameters on the error in thickness measurement (Figure S1E). Those results defined a region in which the predicted error is below 30 nm, which is less than 20% of the typical experimental thickness. We thus used this simulation result as a criterion for image quality, and discarded images or regions of images where the predicted error is out of this accepted region (Figure S1E).

##### TEM and CLEM images

The full cell wall in electron microscopy images was detected using custom scripts written in MATLAB, by using a threshold value for the signal intensity. In some images with inhomogeneous contrast, the results were manually updated. The edge of the binarized signal were traced to define the inside and outside boundaries of the cell wall. The normal direction was defined based on the inside cell wall boundary, and the cell wall thickness was defined as the distance between inside and outside boundaries along the normal direction. To compare EM and light microscopy images in CLEM analyses (Figure 1B first panel and S2E-G), we defined four morphologically distinguishable points (two tips and two scars), and used these points to overlay the two images by matching the distance between them. Comparison on larger region, was done by measuring thickness in EM pictures manually within 2µm regions at 8 individual positions, and compared with an averaged computed from light-microscopy measurement in the Matlab script (Figure 1B third panel).

##### Cell shape, growth and polarity factor concentration measurements

Cell length was generally measured using lectin signals in ImageJ. To compute old end elongation rates visible birth scars were used as fiducial reference landmarks. Tip curvature was computed from the lectin contour defined for cell wall measurements, by fitting a local circle in Matlab (Bonazzi et al., 2015).

To analyze changes in cell shape, growth rates and localization of polar factors following turgor reduction by sorbitol addition, we developed dedicated Matlab scripts. We first segmented cells using the signal from the lectin-labelled cell wall. To this aim, we first smoothed the image with median and Gaussian filters and detected cell edges using the Laplacian of the Gaussian filter. The resultant binary image was then filtered to remove small segments. Given that the signal of the labelled cell wall has a finite thickness, we detected the inner and the outer border of this signal. All spaces in this image are then filled in white except for the spaces between the inner and outer border of the wall, yielding a black band representing the cell wall. Using the watershed algorithm, we finally extracted the whole-cell contour defined as the middle of this band. To compute cell length, we fitted the long axis of the segmented cell with a polynomial of degree 3. This fit was then used to define a “cell spine” and its length was calculated, and used as a measure for cell length.

The whole-cell contour could then be manipulated using morphological and logical operations to obtain a set of arbitrary regions (tips, cytoplasm, etc.). The tip regions are for instance shaped as a cut off from the whole-cell mask perpendicular to the cell spine at specific distances along the spine. A segment outside of the cell can be shaped to compute the background.

Fluorescent signals of interest are then extracted from fluorescent images by using a mask based on corresponding sub-regions. The signal from the tips is corrected for the background signal and for bleaching. This is done by normalizing the background corrected tip signal with the background corrected cytoplasm signal:

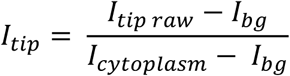

#### Strain-stress assay simulation

The estimation procedure of the mechanical parameters is similar to the one described in the paragraph “Measurement of material properties” in Abenza-Martinez *et al.* (Abenza et al., 2015) except herein the cell wall thickness has been measured all around the contour. For each of the 21 cells, the steps are:

1. Symmetrization of both the plasmolized contour and the turgid contour of each cell; the same symmetrization is kept for the thickness profile.
2. The plasmolyzed contour is swelled at different pressure steps (pressure is normalized by the Young modulus at the equator) supposing a distribution of thickness, and a distribution of mechanical parameters away from the equator; two different distributions of thickness were tested: averaged plasmolyzed thickness and measured plasmolyzed thickness; three distributions of Young modulus were tested: a zero-parameter (i.e constant Young modulus) and two two-parameters distributions (“cos” and “sigmoid”), one parameter describing the tip width, the other parameter being the ratio between the Young modulus at the tip and the Young modulus at the equator. As two recent articles (Abenza et al., 2015; Atilgan et al., 2015) consistently reported a zero-value of the in-plane Poisson ratio, the in-plane Poisson ratio was settled to 0.01 for all the simulations.
3. Among the pressure steps and the tested parameters, the ones for which the swelled plasmolyzed contour best fit the turgid contour are retained to estimate the mechanical parameters. Whatever is the chosen Young modulus distribution, the selected normalized pressure step provides the ratio between the Young modulus at the equator and the internal turgor. The turgor pressure inside the cell has been estimated to be 1.5 MPa in Atilgan *(Atilgan et al., 2015)* providing an absolute estimate of the Young moduli.

Variation in local Young moduli where simulated using:

a) A “sigmoid” distribution on the form: Y(s)=Y_tip_+(Y_side_-Y_tip_)(sigmf(s,[5 d])-sigmf(s,[5 arc-d]))). With s, the curvilinear coordinate and d the width along cell tip. The “sigmoid” distribution is flat at the equator, decreases sharply at a distance d from the tips, then is flat again at the tips.

Deviation from the compuatiaonnaly swollen cell and the orginal cell before turgor reduction was computed as a distance from the best-fit. For tip size of 2.5μm; 3 μm and 3.5μm we obtained respective values for Y_side_/P of 48.1±0.7; 49.0±0.8 and 48.9±0.8; for Y_tip_/Y_side_ = 0.60±0.02; 0.63±0.02 and 0.68±0.02 with distances to best fits of 0.049±0.001; 0.050±0.001; 0.053±0.001.

Those values suggest that the lowest distance from the best fit are obtained for tip-width d small enough. The value of E_cyl_/P are not very sensitive to d whereas E_tip_/E_cyl_ increases with d. The correlation between E_tip_ and E_cyl_ is not significant for d below 3 (p-value>0.5 for both the Pearson test and the Spearman test) whereas it becomes significant for d=3.5 (p-value 0.02 Pearson, 0.11 Spearman) which means the tip width has been chosen too large and that the estimations of both the tip and the equatorial Young modulus are combined in the measurements.

b) A “cos” distribution: Y(s)=Y_tip_ +(Y_cyl_-Y_tip_)(0.5(1-cos(2πs/arc)))^**d**^. The “cos” distribution is narrower around the equator than the sigmoid one especialy for d>1. Similarly to the sigmoid distribution, the best fit are obtained when the Young modulus decreases in the more slowest manner from the equator (d=0.5, d=1). The estimations of Y_cyl_/P are very sensitive to d as the decrease from the equator becomes very sharp when d=2, they also are higher than with the "sigmoid" fit in order to compensate the quicker decrease at the equator. For the best fit (d=0.5, d=1), the estimated ratio of Y_tip_/Yside are slightly lower than the “sigmoid” ones. Obtained values are for exponent d of 0.5; 1 and 2 respectively: Yside/P: 48.6±0.5; 54.9±0.5 and 73.6±0; Ytip/Ycyl: 0.46±0.02; 0.49±0.07 and 0.39 with distances from best-fit of 0.045; 0.048 and 0.065

#### Analytical Models for tip thickness oscillations

Eq (4) of the main text can be written in the form:

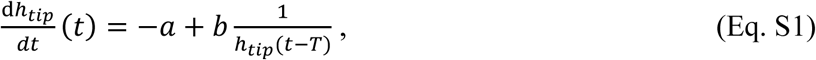

with 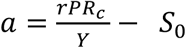, and 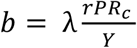.

Using T as a unit of time and the homeostatic thickness value b/a as a unit of thickness, this equation can be rewritten as:

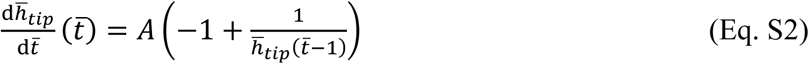

with 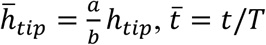, and *A* = *a*^2^*T*/*b*. Numerical solutions of this ordinary differential equation were obtained using the NDSolve function of Mathematica (Wolfram Research Inc.). Eq. S2 has damped solutions with no oscillations for *A* < *A*_1_, oscillating damped solutions for *A*_1_ < *A* < *A*_2_, and sustained oscillations for *A* > *A*_2_. The values of *A*_1_ and *A*_2_ increase when 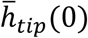 departs from 1. Analytical calculations show that when 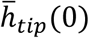 is close to 1, *A*_1_ = exp(1) and *A*_2_ = *π*/2. Optimal homeostasis corresponds to maximal damping, which occurs for, *A* = *A*_1_. Coming back to initial model parameters, 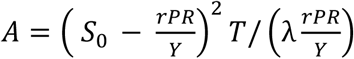. Therefore, a decrease in λ yields an increase in *A*, and hence to oscillations with higher amplitude.

#### 3D cell growth model

The growth model developed in Abenza-Martinez *et al* (Abenza et al., 2015)was modified to include varying thickness and varying young modulus. The parameters h (thickness) and Y (Young modulus) (Formula 36-38 in the supplemental material of (Abenza et al., 2015)) are no more constant.

The thickness distribution shape has been selected by the following procedure: all the symmetrized thickness profiles were linearly reparametrized from new end to old end so that the new end lays at curvilinear abscissa 0 and the old end at a curvilinear abscissa of 13.9. After this operation all the thickness profiles were averaged, the average was very well fitted by a three-parameters linear combination of sigmoids. The three parameters h_NE_, h_OE_ and h_side_ are the cell wall thickness at the new end, at the old end and at the equator.

Only the modification to the original algorithm are detailed here (Supplemental material of Abenza-Martinez *et al* (Abenza et al., 2015)):

- At step *n* (iii), once the new curvilinear abscissa *s*_*n*_ has been calculated using formula 46, *arc*_*n*_, the total arclength is updated and the thickness profile can also be updated using the shape family

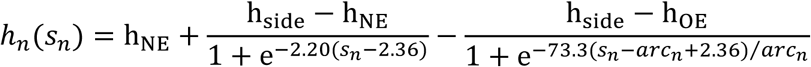

as well as the Young modulus distribution:

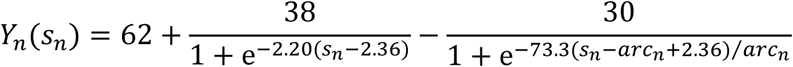

- The amplitude of each step of the growth in formula (47-48) is: δ=1.2

- The model for the fluorescence profile K of the polarity marker:

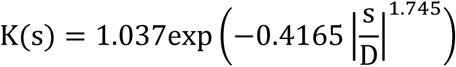

where D=1.17, 1.06 and 0.93 depending on which strain is modeled *rga4Δ*, wt and *rga2Δ*. Each of the D constants have been chosen such that the asymptotic radius of convergence of the cell corresponds to a value close to the experimental tip radii 1.76 μm (*rga4Δ*), 1.62 μm (wt) and 1.46 μm (*rga2Δ*). The cell wall tip thicknesses of *rga4Δ*, wt and *rga2Δ* are 0.149 μm, 0.166 μm and 0.179 μm.

KEY RESOURCES TABLE

KEY RESOURCES TABLE
| REAGENT or RESOURCE | SOURCE | IDENTIFIER |
| --- | --- | --- |
| Experimental Models: *S. pombe* Strains |  |  |
| h- *cut11-RFP:HygR GFP-psy1:leu (leu1-32 ura4-D18)* | This study | VD11 |
| h- *CRIB-tdTomato:ura GFP-psy1:leu (leu1-32 ura4-D18)* | This study | VD20 |
| h- *tea1::NatMX CRIB-tdTomato:ura GFP-psy1:leu (leu1-32 ura4-D18)* | This study | VD31 |
| h+ *CRIB-tdTomato:ura GFP-psy1:leu (leu1-32 ura4-D18)* | This study | VD39 |
| h- *CRIB-tdTomato:ura GFP-psy1:leu orb6-25 (leu1-32 ura4-D18)* | This study | VD50 |
| h+ *GFP-psy1:ade bgs4::ura4 RFP-bgs4-Leu (leu1-32 ura4-D18 ade6)* | This study | VD57 |
| h- *GFP-psy1:ade bgs4::ura4 RFP-bgs4-Leu Leu (leu1-32 ura4-D18 ade6)* | This study | VD58 |
| h- *rga2::KanMX bgs4::ura4 RFP-bgs4-Leu GFP-psy1:ade (leu1-32 ura4-D18)* | This study | VD72 |
| h- *rga4::KanMX bgs4::ura4 RFP-bgs4-Leu GFP-psy1:ade (leu1-32 ura4-D18)* | This study | VD81 |
| h- *GFP-psy1:ade pREP3X-pck2 (leu1-32 ade6)* | This study | VD93 |
| h- *GFP-psy1:ade pRep3X empty vector (leu1-32 ade6)* | This study | VD105 |
| h- *tea1::NatMX GFP-psy1:ade bgs4::ura4 RFP-bgs4:leu (leu1-32 ura4-D18 ade6)* | This study | VD110 |
| *cwg1-1 GFP-psy1:leu CRIB-tdTomato:ura (leu1-32 ura4-D18)* | This study | VD115 |
| *wsc1::KanMX GFP-psy1:ade bgs4::ura4 RFP-bgs4:leu (leu1-32 ura4-D18 ade6)* | This study | VD158 |
| *mtl2::KanMX GFP-psy1:ade bgs4::ura4 RFP-bgs4:leu (leu1-32 ura4-D18 ade6)* | This study | VD160 |
| h<sup>90</sup> *rga4::KanMX GFP-psy1:ade bgs4::ura4 RFP-bgs4:leu (leu1-32 ura4-D18 ade6)* | This study | VD162 |
| *rga4::KanMX GFP-psy1:ade bgs4::ura4 RFP-bgs4:leu (leu1-32 ura4-D18 ade6)* | This study | VD163 |
| *rgf2::ura GFP-psy1:ade (leu1-32 ura4-D18 ade6)* | This study | VD164 |
| h+ *rgf1::KanMX GFP-psy1:ade (leu1-32 ura4-D18 ade6)* | This study | VD72 |
| *pmk1:ura GFP-psy1:leu (leu1-32 ura4-D18)* | This study | VD81 |
| *rgf1::his3 leu1:rgf1-GFP (his3D1 leu1-32 ade6M210 ura4D-18)* | Y. Sanchez* | VD93 |
| h+ *pRep3X-Phluorin-ritc (leu1-32)* | This study | VD105 |
| h+ *Leu:GFP-psy1 (leu1-32 ura4-D18)* | This study | NM468 |
| h- *Leu:GFP-psy1 (leu1-32 ura4-D18)* | This study | NM469 |
| h+ *shk1:GFP-LactC2:leu (leu1-32)* | This study | AH159 |
| *gpd1::NatMX GFP-bgs4:leu bgs4::ura (leu1-32 ura4-D18 ade6)* | This study | AH328 |
| *gpd1::NatMX wsc1::KanMX GFP-bgs4:leu bgs4::ura (ade6)* | This study | AH330 |
| h- *gpd1::NatMX P3nmt1-pkc1(HR1-C2)-ECitrine (ura4-D18)* | This study | AH385 |
| *gpd1::NatMX wsc1::KanMX P3nmt1-pkc1(HR1-C2)-ECitrine (leu1-32 ura4-D18 ade6)* | This study | AH401 |
| *cdc10-M17 GFP-psy1:ade bgs4::ura4 RFP-bgs4-Leu (ade6)* | This study | VD122 |
| *cdc25-22 GFP-psy1:ade bgs4::ura4 RFP-bgs4-Leu (leu1-32 ura4-D18 ade6)* | This study | VD123 |
| Chemicals |  |  |
| Zymolyase | ZymoResearch | E1004 |
| LatrunculinA | Sigma-Aldrich | L5163 |
| Caspofungin | Sigma-Aldrich | SML0425 |
| *Gs*-IB<sub>4</sub> -Alexafluor647 | ThermoFisher | I32450 |
| *Bs*-IB<sub>4</sub>-TRIC | Sigma-Aldrich | N/A |

## REFERENCES

Abenza, J.F., Couturier, E., Dodgson, J., Dickmann, J., Chessel, A., Dumais, J., and Carazo Salas, R.E. (2015). Wall mechanics and exocytosis define the shape of growth domains in fission yeast. Nat Commun 6, 8400.

Arellano, M., Duran, A., and Perez, P. (1996). Rho 1 GTPase activates the (1-3)beta-D-glucan synthase and is involved in Schizosaccharomyces pombe morphogenesis. EMBO J 15, 4584–4591.

Arellano, M., Valdivieso, M.H., Calonge, T.M., Coll, P.M., Duran, A., and Perez, P. (1999). Schizosaccharomyces pombe protein kinase C homologues, pck1p and pck2p, are targets of rho1p and rho2p and differentially regulate cell integrity. J Cell Sci 112 (Pt 20), 3569–3578.

Atilgan, E., Magidson, V., Khodjakov, A., and Chang, F. (2015). Morphogenesis of the Fission Yeast Cell through Cell Wall Expansion. Curr Biol 25, 2150–2157.

Bastmeyer, M., Deising, H.B., and Bechinger, C. (2002). Force exertion in fungal infection. Annu Rev Biophys Biomol Struct 31, 321–341.

Bendezu, F.O., and Martin, S.G. (2011). Actin cables and the exocyst form two independent morphogenesis pathways in the fission yeast. Mol Biol Cell 22, 44–53.

Bonazzi, D., Haupt, A., Tanimoto, H., Delacour, D., Salort, D., and Minc, N. (2015). Actin-Based Transport Adapts Polarity Domain Size to Local Cellular Curvature. Curr Biol 25, 2677–2683.

Bonazzi, D., Julien, J.D., Romao, M., Seddiki, R., Piel, M., Boudaoud, A., and Minc, N. (2014). Symmetry Breaking in Spore Germination Relies on an Interplay between Polar Cap Stability and Spore Wall Mechanics. Dev Cell 28, 534–546.

Boudaoud, A. (2003). Growth of walled cells: from shells to vesicles. Phys Rev Lett 91, 018104.

Cassone, A., Kerridge, D., and Gale, E.F. (1979). Ultrastructural changes in the cell wall of Candida albicans following cessation of growth and their possible relationship to the development of polyene resistance. J Gen Microbiol 110, 339–349.

Chang, F., and Huang, K.C. (2014). How and why cells grow as rods. BMC Biol 12, 54.

Chang, F., and Martin, S.G. (2009). Shaping fission yeast with microtubules. Cold Spring Harb Perspect Biol 1, a001347.

Charvin, G., Cross, F.R., and Siggia, E.D. (2008). A microfluidic device for temporally controlled gene expression and long-term fluorescent imaging in unperturbed dividing yeast cells. PLoS One 3, e1468.

Chugh, P., Clark, A.G., Smith, M.B., Cassani, D.A.D., Dierkes, K., Ragab, A., Roux, P.P., Charras, G., Salbreux, G., and Paluch, E.K. (2017). Actin cortex architecture regulates cell surface tension. Nat Cell Biol 19, 689–697.

Clark, A.G., Dierkes, K., and Paluch, E.K. (2013). Monitoring actin cortex thickness in live cells. Biophys J 105, 570–580.

Cortes, J.C., Carnero, E., Ishiguro, J., Sanchez, Y., Duran, A., and Ribas, J.C. (2005). The novel fission yeast (1,3)beta-D-glucan synthase catalytic subunit Bgs4p is essential during both cytokinesis and polarized growth. J Cell Sci 118, 157–174.

Cosgrove, D.J. (2005). Growth of the plant cell wall. Nat Rev Mol Cell Biol 6, 850–861.

Cruz, S., Munoz, S., Manjon, E., Garcia, P., and Sanchez, Y. (2013). The fission yeast cell wall stress sensor-like proteins Mtl2 and Wsc1 act by turning on the GTPase Rho1p but act independently of the cell wall integrity pathway. Microbiologyopen 2, 778–794.

Dagdas, Y.F., Yoshino, K., Dagdas, G., Ryder, L.S., Bielska, E., Steinberg, G., and Talbot, N.J. (2012). Septin-mediated plant cell invasion by the rice blast fungus, Magnaporthe oryzae. Science 336, 1590–1595.

Das, M., Wiley, D.J., Chen, X., Shah, K., and Verde, F. (2009). The conserved NDR kinase Orb6 controls polarized cell growth by spatial regulation of the small GTPase Cdc42. Curr Biol 19, 1314–1319.

Das, M., Wiley, D.J., Medina, S., Vincent, H.A., Larrea, M., Oriolo, A., and Verde, F. (2007). Regulation of cell diameter, For3p localization, and cell symmetry by fission yeast Rho-GAP Rga4p. Mol Biol Cell 18, 2090–2101.

Davi, V., and Minc, N. (2015). Mechanics and morphogenesis of fission yeast cells. Curr Opin Microbiol 28, 36–45.

Davidson, R., Laporte, D., and Wu, J.Q. (2015). Regulation of Rho-GEF Rgf3 by the arrestin Art1 in fission yeast cytokinesis. Mol Biol Cell 26, 453–466.

DeBerardinis, R.J., Lum, J.J., Hatzivassiliou, G., and Thompson, C.B. (2008a). The biology of cancer: metabolic reprogramming fuels cell growth and proliferation. Cell Metab 7, 11–20.

Deberardinis, R.J., Sayed, N., Ditsworth, D., and Thompson, C.B. (2008b). Brick by brick: metabolism and tumor cell growth. Curr Opin Genet Dev 18, 54–61.

Drake, T., and Vavylonis, D. (2013). Model of fission yeast cell shape driven by membrane-bound growth factors and the cytoskeleton. PLoS Comput Biol 9, e1003287.

Dudin, O., Bendezu, F.O., Groux, R., Laroche, T., Seitz, A., and Martin, S.G. (2015). A formin-nucleated actin aster concentrates cell wall hydrolases for cell fusion in fission yeast. J Cell Biol 208, 897–911.

DuFort, C.C., Paszek, M.J., and Weaver, V.M. (2011). Balancing forces: architectural control of mechanotransduction. Nat Rev Mol Cell Biol 12, 308–319.

Fantes, P., and Nurse, P. (1977). Control of cell size at division in fission yeast by a growth-modulated size control over nuclear division. Exp Cell Res 107, 377–386.

Fink, J., Carpi, N., Betz, T., Betard, A., Chebah, M., Azioune, A., Bornens, M., Sykes, C., Fetler, L., Cuvelier, D., et al. (2011). External forces control mitotic spindle positioning. Nat Cell Biol 13, 771–778.

Flor-Parra, I., Zhurinsky, J., Bernal, M., Gallardo, P., and Daga, R.R. (2014). A Lallzyme MMX-based rapid method for fission yeast protoplast preparation. Yeast 31, 61–66.

Garcia, P., Garcia, I., Marcos, F., de Garibay, G.R., and Sanchez, Y. (2009a). Fission yeast rgf2p is a rho1p guanine nucleotide exchange factor required for spore wall maturation and for the maintenance of cell integrity in the absence of rgf1p. Genetics 181, 1321–1334.

Garcia, P., Tajadura, V., Garcia, I., and Sanchez, Y. (2006). Rgf1p is a specific Rho1-GEF that coordinates cell polarization with cell wall biogenesis in fission yeast. Mol Biol Cell 17, 1620–1631.

Garcia, P., Tajadura, V., and Sanchez, Y. (2009b). The Rho1p exchange factor Rgf1p signals upstream from the Pmk1 mitogen-activated protein kinase pathway in fission yeast. Mol Biol Cell 20, 721–731.

Harold, F.M. (2002). Force and compliance: rethinking morphogenesis in walled cells. Fungal Genet Biol 37, 271–282.

Harris, L.K., and Theriot, J.A. (2016). Relative Rates of Surface and Volume Synthesis Set Bacterial Cell Size. Cell 165, 1479–1492.

Heinisch, J.J., Dupres, V., Wilk, S., Jendretzki, A., and Dufrene, Y.F. (2010). Single-molecule atomic force microscopy reveals clustering of the yeast plasma-membrane sensor Wsc1. PLoS One 5, e11104.

Hepler, P.K., Vidali, L., and Cheung, A.Y. (2001). Polarized cell growth in higher plants. Annu Rev Cell Dev Biol 17, 159–187.

Holley, R.W. (1975). Control of growth of mammalian cells in cell culture. Nature 258, 487–490.

Hong, L., Dumond, M., Tsugawa, S., Sapala, A., Routier-Kierzkowska, A.L., Zhou, Y., Chen, C., Kiss, A., Zhu, M., Hamant, O., et al. (2016). Variable Cell Growth Yields Reproducible OrganDevelopment through Spatiotemporal Averaging. Dev Cell 38, 15–32.

Horiseberger, M., and Rosset, J. (1977). Localization of alpha-Galactomannan on the surface of Schizosaccharomyces pombe cells by scanning electron microscopy. Arch Microbiol 112, 123–126.

Houk, A.R., Jilkine, A., Mejean, C.O., Boltyanskiy, R., Dufresne, E.R., Angenent, S.B., Altschuler, S.J., Wu, L.F., and Weiner, O.D. (2012). Membrane tension maintains cell polarity by confining signals to the leading edge during neutrophil migration. Cell 148, 175–188.

Howell, A.S., Jin, M., Wu, C.F., Zyla, T.R., Elston, T.C., and Lew, D.J. (2012). Negative feedback enhances robustness in the yeast polarity establishment circuit. Cell 149, 322–333.

Huang, S., and Ingber, D.E. (1999). The structural and mechanical complexity of cell-growth control. Nat Cell Biol 1, E131–138.

Keegstra, K. (2010). Plant cell walls. Plant Physiol 154, 483–486.

Kelly, F.D., and Nurse, P. (2011). De novo growth zone formation from fission yeast spheroplasts. PLoS One 6, e27977.

Kierzkowski, D., Nakayama, N., Routier-Kierzkowska, A.L., Weber, A., Bayer, E., Schorderet, M., Reinhardt, D., Kuhlemeier, C., and Smith, R.S. (2012). Elastic domains regulate growth and organogenesis in the plant shoot apical meristem. Science 335, 1096–1099.

Lander, A.D. (2011). Pattern, growth, and control. Cell 144, 955–969.

Levayer, R., and Lecuit, T. (2012). Biomechanical regulation of contractility: spatial control and dynamics. Trends Cell Biol 22, 61–81.

Lew, R.R. (2011). How does a hypha grow? The biophysics of pressurized growth in fungi. Nat Rev Microbiol 9, 509–518.

Lipke, P.N., and Ovalle, R. (1998). Cell wall architecture in yeast: new structure and new challenges. J Bacteriol 180, 3735–3740.

Ma, H., Snook, L.A., Kaminskyj, S.G., and Dahms, T.E. (2005). Surface ultrastructure and elasticity in growing tips and mature regions of Aspergillus hyphae describe wall maturation. Microbiology 151, 3679–3688.

Mahadevan, L., and Mitchison, T.J. (2005). Cell biology: powerful curves. Nature 435, 895–897.

Marshall, W.F., Young, K.D., Swaffer, M., Wood, E., Nurse, P., Kimura, A., Frankel, J., Wallingford, J., Walbot, V., Qu, X., et al. (2012). What determines cell size? BMC Biol 10, 101.

Martins, I.M., Cortes, J.C., Munoz, J., Moreno, M.B., Ramos, M., Clemente-Ramos, J.A., Duran, A., and Ribas, J.C. (2011). Differential activities of three families of specific beta(1,3)glucan synthase inhibitors in wild-type and resistant strains of fission yeast. J Biol Chem 286, 3484–3496.

Mata, J., and Nurse, P. (1997). tea1 and the microtubular cytoskeleton are important for generating global spatial order within the fission yeast cell. Cell 89, 939–949.

McKenna, S.T., Kunkel, J.G., Bosch, M., Rounds, C.M., Vidali, L., Winship, L.J., and Hepler, P.K. (2009). Exocytosis precedes and predicts the increase in growth in oscillating pollen tubes. Plant Cell 21, 3026–3040.

Miermont, A., Uhlendorf, J., McClean, M., and Hersen, P. (2011). The Dynamical Systems Properties of the HOG Signaling Cascade. J Signal Transduct 2011, 930940.

Milani, P., Mirabet, V., Cellier, C., Rozier, F., Hamant, O., Das, P., and Boudaoud, A. (2014). Matching Patterns of Gene Expression to Mechanical Stiffness at Cell Resolution through Quantitative Tandem Epifluorescence and Nanoindentation. Plant Physiol 165, 1399–1408.

Minc, N., Boudaoud, A., and Chang, F. (2009). Mechanical forces of fission yeast growth. Curr Biol 19, 1096–1101.

Mitchison, J.M., and Nurse, P. (1985). Growth in cell length in the fission yeast Schizosaccharomyces pombe. J Cell Sci 75, 357–376.

Mogilner, A., and Keren, K. (2009). The shape of motile cells. Curr Biol 19, R762–771.

Munoz, J., Cortes, J.C., Sipiczki, M., Ramos, M., Clemente-Ramos, J.A., Moreno, M.B., Martins, I.M., Perez, P., and Ribas, J.C. (2013). Extracellular cell wall beta(1,3)glucan is required to couple septation to actomyosin ring contraction. J Cell Biol 203, 265–282.

Murray, S. (2008). High pressure freezing and freeze substitution of Schizosaccharomyces pombe and Saccharomyces cerevisiae for TEM. Methods Cell Biol 88, 3–17.

Mutoh, T., Nakano, K., and Mabuchi, I. (2005). Rho1-GEFs Rgf1 and Rgf2 are involved in formation of cell wall and septum, while Rgf3 is involved in cytokinesis in fission yeast. Genes Cells 10, 1189–1202.

Nakamura, T., Nakamura-Kubo, M., Hirata, A., and Shimoda, C. (2001). The Schizosaccharomyces pombe spo3+ gene is required for assembly of the forespore membrane and genetically interacts with psy1(+)-encoding syntaxin-like protein. Mol Biol Cell 12, 3955–3972.

Nakayama, N., Smith, R.S., Mandel, T., Robinson, S., Kimura, S., Boudaoud, A., and Kuhlemeier, C. (2012). Mechanical regulation of auxin-mediated growth. Curr Biol 22, 1468–1476.

Novick, P., and Schekman, R. (1979). Secretion and cell-surface growth are blocked in a temperature-sensitive mutant of Saccharomyces cerevisiae. Proc Natl Acad Sci U S A 76, 1858–1862.

Osumi, M. (2012). Visualization of yeast cells by electron microscopy. J Electron Microsc (Tokyo) 61, 343–365.

Peaucelle, A., Braybrook, S.A., Le Guillou, L., Bron, E., Kuhlemeier, C., and Hofte, H. (2011). Pectin-induced changes in cell wall mechanics underlie organ initiation in Arabidopsis. Curr Biol 21, 1720–1726.

Perez, P., and Cansado, J. (2010). Cell integrity signaling and response to stress in fission yeast. Curr Protein Pept Sci 11, 680–692.

Perez, P., and Ribas, J.C. (2004). Cell wall analysis. Methods 33, 245–251.

Petersen, J., Heitz, M.J., and Hagan, I.M. (1998). Conjugation in S. pombe: identification of a microtubule-organising centre, a requirement for microtubules and a role for Mad2. Curr Biol 8, 963–966.

Qin, Y., and Yang, Z. (2011). Rapid tip growth: insights from pollen tubes. Semin Cell Dev Biol 22, 816–824.

Rojas, E., Theriot, J.A., and Huang, K.C. (2014). Response of Escherichia coli growth rate to osmotic shock. Proc Natl Acad Sci U S A 111, 7807–7812.

Rojas, E.R., Hotton, S., and Dumais, J. (2011). Chemically mediated mechanical expansion of the pollen tube cell wall. Biophys J 101, 1844–1853.

Salbreux, G., Charras, G., and Paluch, E. (2012). Actin cortex mechanics and cellular morphogenesis. Trends Cell Biol 22, 536–545.

Silhavy, T.J., Kahne, D., and Walker, S. (2010). The bacterial cell envelope. Cold Spring Harb Perspect Biol 2, a000414.

Tatebe, H., Nakano, K., Maximo, R., and Shiozaki, K. (2008). Pom1 DYRK regulates localization of the Rga4 GAP to ensure bipolar activation of Cdc42 in fission yeast. Curr Biol 18, 322–330.

Thompson, B.J. (2010). Developmental control of cell growth and division in Drosophila. Curr Opin Cell Biol 22, 788–794.

Villar-Tajadura, M.A., Coll, P.M., Madrid, M., Cansado, J., Santos, B., and Perez, P. (2008). Rga2 is a Rho2 GAP that regulates morphogenesis and cell integrity in S. pombe. Mol Microbiol 70, 867–881.

Yanagisawa, M., Desyatova, A.S., Belteton, S.A., Mallery, E.L., Turner, J.A., and Szymanski, D.B. (2015). Patterning mechanisms of cytoskeletal and cell wall systems during leaf trichome morphogenesis. Nat Plants 1, 15014.

Zegman, Y., Bonazzi, D., and Minc, N. (2015). Measurement and manipulation of cell size parameters in fission yeast. In Methods in Cell Biology (Academic Press).

